# High-resolution population structure inference using genome-wide short tandem repeat variations

**DOI:** 10.64898/2026.02.20.707006

**Authors:** Feifei Xia, Michael Baudis, Maria Anisimova

## Abstract

Short tandem repeats (STRs) are a major source of genetic variation, yet their potential for genome-wide population structure inference remains underexplored. Here we present a multi-modal framework for STR-based population inference, integrating unsupervised clustering, supervised population assignment, and a novel admixture inference model, Directional Non-negative Matrix Factorization (dNMF). Applying this framework to thousands of genomes from multiple global cohorts, we first demonstrate that genome-wide STR variations provide substantially finer resolution of human population structure than single-nucleotide polymorphisms (SNPs), particularly at regional levels. The dNMF model estimates ancestry coefficients under the hypothesis that true ancestral populations are consistently encoded in the bidirectional mutation dynamics of STRs. Population structures inferred by dNMF are fine-grained, reproducible across datasets, and robust to technical artifacts. Motif-specific analyses further reveal direction-biased mutational tendencies and show that distinct STR motif classes encode complementary layers of population structure at different evolutionary scales. These results establish STRs as powerful and biologically interpretable markers for population structure inference, offering a mutation-aware perspective that complements traditional SNP-based frameworks and refines understanding of human demographic history.

## Introduction

Understanding human population structure and genetic diversity is central to evolutionary biology and population genetics. Over the past decades, large-scale analyses of population structure have been dominated by inferences from single-nucleotide polymorphisms (SNPs), which are abundant, stable, and easily assayed at scale. SNP-based approaches, such as principal component analysis (PCA) and model-based inference of ancestry proportions^1–5^, have provided deep insights into human demographic history, migration, and genetic diversity. Historically, short tandem repeats (STRs), also known as microsatellites, were among the first molecular markers used to characterize human population structure and genetic diversity^6–10^. Their high polymorphism and multi-allelic nature made them ideal for early studies of population differentiation, migration, and relatedness. However, as SNP genotyping technology became more efficient and scalable, SNP-based analyses gradually replaced STRs in large-scale studies of population structure inference^11–13^. Consequently, STRs have remained largely underutilized at the genome-wide scale, despite their wide application in forensic analysis and paternity testing.

STRs consist of repeat units of 1-6 bp and represent one of the largest sources of human genetic variation^14,15^. With their high mutation rates and multi-allelic nature, STRs hold considerable potential for resolving recent demographic events and subtle population differentiation^16,17^. Despite their promise, STRs have not yet been systematically applied to genome-wide population structure inference. Several studies have performed genome-wide analyses of STR variations in human population^18–22^, including comprehensive catalogs of STR polymorphisms across global populations^23,24^. However, these efforts primarily characterized STR diversity, mutation patterns, and population allele frequency distributions rather than explicitly inferring population structure.

Recent advances in high-coverage sequencing and specialized STR genotyping algorithms^25–27^, have revitalized interest in STRs as genome-wide markers across human populations. Moreover, STR-based genome-wide association studies have revealed functional roles of STR variation in gene regulation and complex traits^28–32^. Together, these studies suggest that STRs capture a complementary layer of genetic variations that remain underexplored in population and functional genomics, with the opportunity to revisit STRs as genetic markers of population structure at the genome-wide level.

Most previous studies have relied on limited forensic panels or small sets of loci, falling short of the comprehensive frameworks developed for SNPs. While methods such as principal component analysis can be applied to quantitative STR genotypes, model-based ancestry inference frameworks, including admixture and related approaches, were originally designed for binary or diploid SNP genotypes and are therefore not directly suited to analyse the multi-allelic and quantitative nature of STR variation. It remains unclear whether genome-wide STRs can recapitulate known patterns of human population structure, whether they provide complementary resolution beyond SNPs, and how robustly STR-derived structures generalize across independent datasets. Addressing these questions requires dedicated framework that can exploit the high variability of STRs, assess their reproducibility, and account for mutational processes that may obscure demographic signals.

To unlock the full potential of STRs, we developed a comprehensive, multi-modal framework for STR-based population inference. This framework integrates three complementary analytical approaches: unsupervised clustering with principal component analysis (PCA) and t-SNE for exploratory data analysis, supervised machine learning models for high-accuracy population assignment, and a novel admixture model, Directional Non-negative Matrix Factorization (dNMF), designed to decouple ancestry structure from underlying STR mutational mechanics. In this study, we apply our framework to thousands of genomes from diverse global cohorts, including the 1000 Genomes Project (1KGP)^33^, Human Genome Diversity Project (HGDP)^34^, Simon Genome Diversity Project (SGDP)^35^, and H3Africa^36^. First, we demonstrate that genome-wide STRs provide higher resolution in regional population inference than SNPs in both unsupervised and supervised analyses, and that the resulting population assignments are robust and reproducible across datasets. We then apply our dNMF model to show that true ancestral structure is consistently encoded across both directions of STR mutation. Finally, we uncover reproducible population-specific signatures of STR mutational dynamics that likely reflect fundamental, genome-wide forces rather than locus-specific selection. Together, this work establishes a new paradigm for STR-based population genetics, presenting a powerful analytical framework and offering novel insights into the mutational processes that shape human genetic diversity.

## Results

### A multi-modal framework for STR-based population structure inference

We obtained genome-wide STR genotypes for 3202 samples from the 1000 Genomes Project (1KGP) and 348 samples from the H3Africa Project, both accessed through the population reference panel^24^. We focused on STRs of 1-6 base pairs and selected genotypes generated by HipSTR^25^, which showed the highest Mendelian inheritance rate in the original study. In addition, we performed genome-wide genotyping of STRs using high-coverage WGS data for 828 samples from the Human Genome Diversity Project (HGDP)^34^ and 276 samples from the Simon Genome Diversity Project (SGDP)^35^. After filtering low-quality calls (Methods), we calculated the mean allele length for each STR locus and retained variable loci to construct STR genotype matrices for downstream analysis (Fig. 1a, Supplementary Data 1).

**Figure 1:**
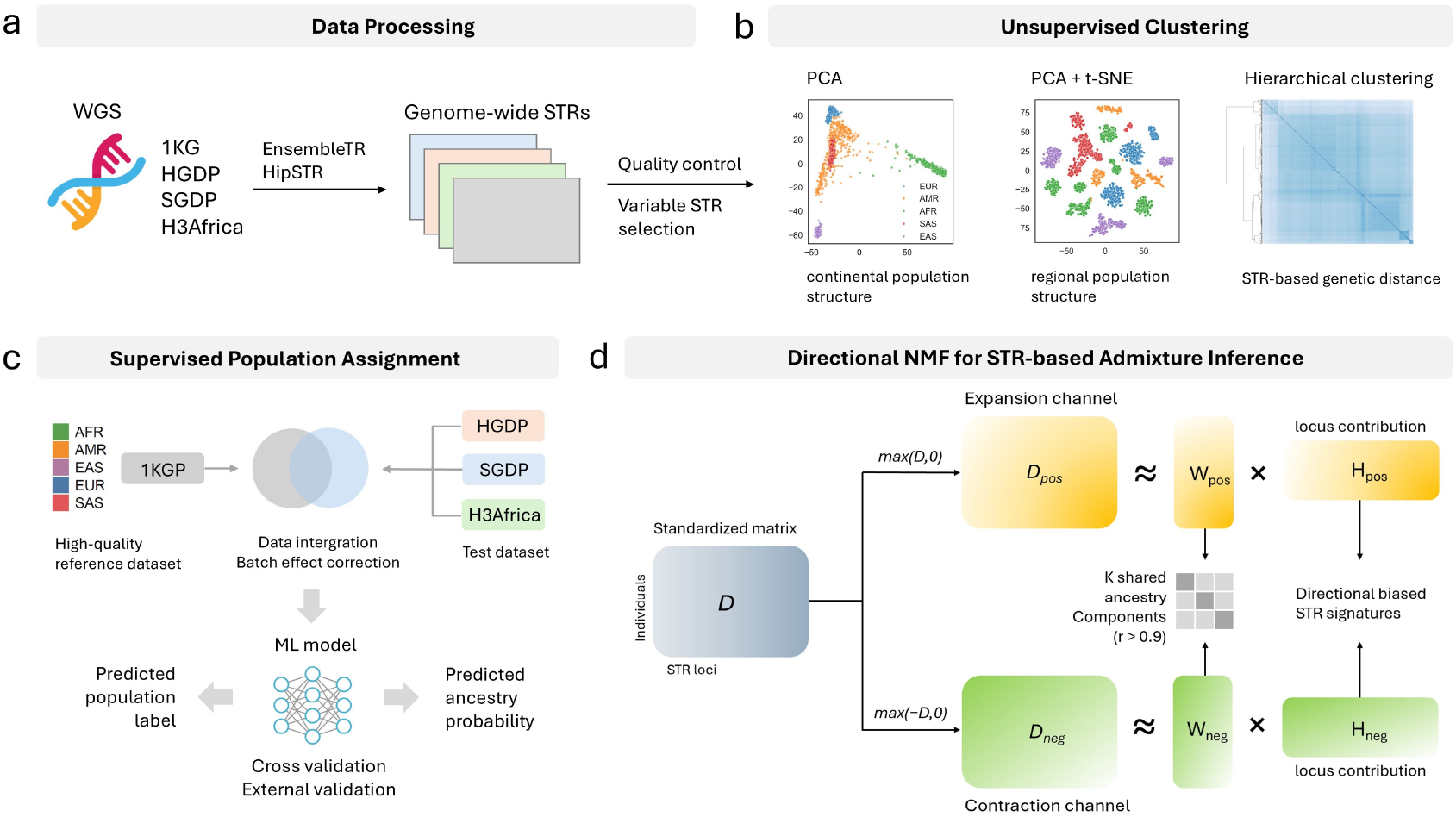
A multi-modal framework for population inference using genome-wide STRs. **a** Data processing. Genome-wide STR genotypes were collected or generated using whole-genome sequencing (WGS) data from global cohorts (1KGP, HGDP, SGDP, and H3Africa). After quality control and selection of variable loci, STR genotype matrices containing mean allele length of variable STRs were prepared for downstream analyses. **b** Unsupervised clustering. Principal component analysis (PCA) and non-linear embedding (t-SNE) were used to identify major axes of genetic variation and visualize continental and regional population structure. STR-based genetic distances were further examined by hierarchical clustering. **c** Supervised population assignment. Machine learning models trained on 1KGP reference data were validated on independent datasets (HGDP, SGDP, H3Africa). Cross-validation and external testing assessed the predictive accuracy of STRs for population classification and ancestry probability estimation. **d** Directional NMF (dNMF) is a two-channel matrix factorization framework designed for STR-based admixture inference. It decomposes the standardized STR deviation matrix into expansion and contraction channels, each of which went through independent non-negative factorizations. The model extracts latent ancestry components for individuals while simultaneously learning locus-specific contributions within each channel. A cross-channel alignment procedure identifies shared ancestry components that are invariant to mutational direction. dNMF outputs (i) ancestry proportion matrices for population structure inference and visualization, and (ii) directionally resolved locus-contribution profiles for motif-level analysis.

To establish a systematic basis for STR-based population structure inference, we developed a comprehensive, multi-modal framework that integrates three complementary analytical perspectives: unsupervised clustering, supervised population assignment, and a novel STR-based admixture inference method termed Directional Non-negative Matrix Factorization (dNMF). In the supervised setting, we applied principal component analyses (PCA), t-Distributed Stochastic Neighbor Embedding (t-SNE), and hierarchical clustering to evaluate the patterns of continental and regional population structure capture by genome-wide STRs (Fig. 1b). To benchmark STR performance, we conducted parallel analyses using genome-wide SNPs from the 1KGP and HGDP datasets. We next assessed the predictive power in a supervised setting using multiple machine learning classifiers (Fig. 1c). Models trained on STRs or SNPs were evaluated at both continental and regional population levels. To assess generalizability, we trained classifiers using 1KGP as a reference panel and validated them on independent cohorts, including HGDP, SGDP, and H3Africa. These analyses provided a comprehensive assessment of the resolution and transferability of STR-based population assignment.

Finally, we introduced Directional Non-negative Matrix Factorization (dNMF), an admixture inference model specifically designed for STR variations (Fig. 1d). Traditional admixture frameworks^9^ assume that an individual’s ancestry is a mixture of contributions from multiple ancestral populations. Under the stepwise mutation model, STR allele lengths mutate through single-unit gains and losses, generating a symmetric mutation process in which allele lengths drift over time. Building on these principles, we further propose that the underlying ancestral structure is simultaneously encoded in both STR expansion and contraction directions, providing the conceptual foundation for the directional formulation of dNMF. To capture the directional mutation process, dNMF takes paired matrices of STR expansions and contractions as input data and fits two independent NMF decompositions to derive the corresponding ancestry components **W**_*pos*_ and **W**_*neg*_. By comparing the directional decompositions, dNMF evaluates the consistency of shared ancestral components across mutation directions and identifies motif-specific biases that reflect distinct mutational mechanisms.

Together, this framework integrates complementary population genetic perspectives and establishes a foundation for evaluating the resolution, reproducibility, and biological insights provided by genome-wide STR variations. In the following sections, we benchmark STR-based against SNP-based approaches across unsupervised and supervised settings and assess the robustness of STR-based population structure inference across diverse global cohorts.

### STRs provide enhanced resolution of human population structure compared with SNPs

In both unsupervised and supervised settings, we compared population structure patterns captured by STRs and SNPs. A total of 31,395 variable STRs and 144,912 polymorphic SNPs from the 1KGP dataset were analyzed. First, PCA was performed on genotype matrices derived from each variant type, followed by unsupervised clustering using the k-means algorithm. Clustering concordance with known population labels was quantified using the Adjusted Rand Index (ARI) at both continental and regional population levels (Fig. 2a). Both STRs and SNPs reached a maximum clustering accuracy of 86% when using the first three principal components (PCs), with accuracy plateauing when additional PCs were included, indicating broad agreement in global ancestry patterns. (Supplementary Fig.S1a). However, STRs revealed additional substructure within continental populations, most prominently among African regional populations (accuracy of 93% for STRs vs 70% for SNPs). This pattern is consistent with the higher STR variation observed in African populations and their greater fine-scale genetic diversity^10,24^. Supporting this observation, t-SNE applied to the first 30 PCs displayed clearer separation of regional populations for STRs than for SNPs (Fig. 2b; Supplementary Fig.S2).

**Figure 2:**
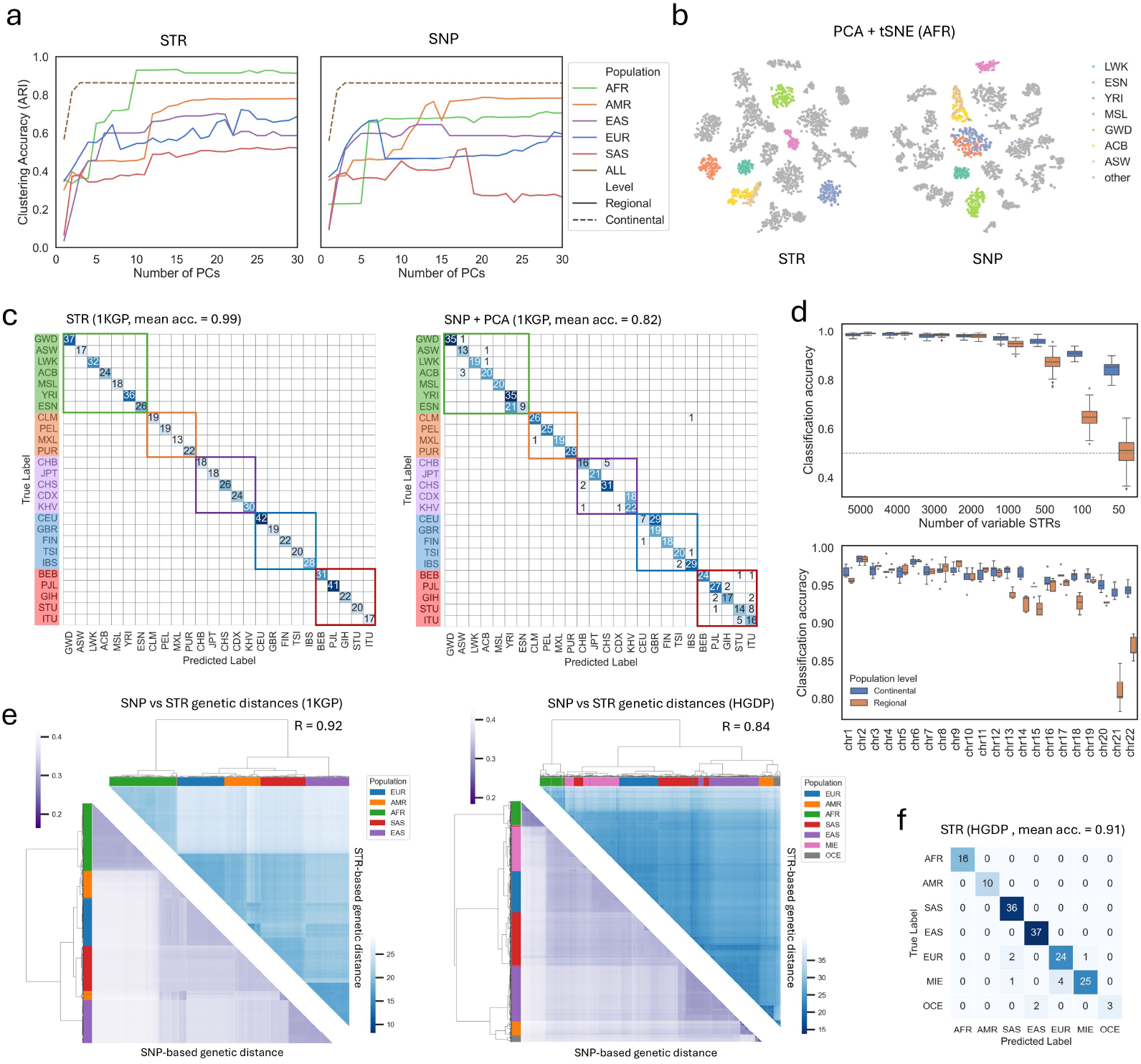
Comparative analysis of STR- and SNP-based population structure inference. **a** Unsupervised clustering accuracy based on STR- and SNP-based principal component analysis (PCA) results across continental and regional population levels in the 1KGP. The x-axis denotes number of principal components (PCs) used for clustering, and the y-axis denotes the k-means clustering accuracy (Adjusted Rand Index, ARI). **b** Two-dimensional visualization of African (AFR) samples using the top 30 PCs followed by t-SNE, showing distinct regional population clusters derived from STRs and SNPs. Samples from non-african populations are shown in grey. **c** Confusion matrices of random forest (RF) regional population assignment in the 1KGP dataset using raw STR genotypes (left) and SNP-derived PCs (right). Mean accuracies were derived from five-fold cross validation. STR-based models achieved markedly higher classification accuracy (mean accuracy = 0.99 vs. 0.82). **d** Continental and regional population assignment accuracies as a function of the number of down-sampled STR loci (top) and per-chromosome STR subsets (bottom). Down-sampling was repeated 10 times per subset, and accuracies were recorded from five-fold cross-validation. **e** Comparison of pairwise genetic distance matrices derived from STRs and SNPs in the 1KGP (left) and HGDP (right) datasets. Heatmaps display STR-based genetic distances (lower triangles) and SNP-based genetic distances (upper triangles), both hierarchically clustered. Each leaf indicate a single individual, with continental population labels shown along the dendrograms. STR-based and SNP-based genetic distance matrices show strong concordance (Pearson’s r = 0.92 and 0.84, respectively. Mantel test, *P <* 10^−3^). **f** Confusion matrix of STR-based continental population assignment in the HGDP dataset. Mean accuracy was calculated from five-fold cross validation.

We next quantified the predictive power of STRs for supervised population assignment using random forest and naïve Bayes classifiers. Random forest models were trained on genotype matrices derived from either STRs or SNPs from the 1KGP dataset and evaluated through stratified five-fold cross-validation at both continental and regional population levels. For SNPs, the models were trained on the first 30 PCs to summarize population structure, whereas STR-based models were trained directly on the raw genotype matrices without dimensionality reduction. Both variant types achieved near-perfect accuracy at the continental level, again indicating strong concordance. However, STR-based models consistently outperformed SNP-based models in distinguishing regional populations, achieving 99% versus 82% accuracy, respectively (Fig. 2c). Notably, SNP-based classification required dimensionality reduction to achieve robust accuracy, whereas STRs achieved near-perfect classification accuracy using raw genotypes, reflecting their higher informativeness per locus. Furthermore, comparable accuracy was maintained when STRs were analyzed with a simple naïve Bayes classifier, confirming that the high predictive power of STRs arises from their intrinsic genetic informativeness rather than from model complexity. Collectively, these results showed that STRs provided greater discriminatory power for recent demographic differentiation compared to biallelic SNPs.

To further assess the robustness and genomic distribution of STR-based ancestry signals, we evaluated how classification accuracy changed with the number of variable STRs included in downsampled marker sets, as well as subsets of variable STRs available on each chromosome (Methods; Fig. 2d). STR-based classification remained near-perfect even when the marker set was reduced to 1,000 loci, suggesting that only a modest number of variable STRs are sufficient to recover global and regional population structure. With further reduction in marker set size, accuracy declined more slowly at the continental level than at the regional level, reflecting the greater amount of information required to resolve fine-scale structure. When evaluated on a per-chromosome basis, most autosomes achieved classification accuracies above 95% at the continental level, with chromosomes 1–16 retaining strong regional resolution. On average, the number of variable STRs per chromosome was 1,425, and accuracy decreased on chromosomes with substantially fewer informative loci, particularly chromosome 22. These results demonstrate that ancestry-informative STRs are broadly distributed across the human genome and that population structure inference remains highly accurate even with substantially reduced marker sets.

Finally, we compared SNP-based and STR-based genetic distances to evaluate the concordance of genetic differentiation inferred from the two variant types. Pairwise genetic distance matrices were computed among individuals from the 1KGP and HGDP datasets using the Goldstein distance^37^ for STRs, which accounts for differences in repeat length under the stepwise mutation model, and the allele-sharing distance for SNPs. Hierarchical clustering of these matrices revealed broadly consistent population relationships across both variant types (Fig. 3e). This concordance was further supported by comparing regional population genetic distances with geographical distances within each continental population, which yielded consistent patterns in both datasets (Supplementary Fig.S3). We additionally performed Mantel tests between the STR- and SNP-based distance matrices, yielding correlation coefficients of 0.92 for 1KGP and 0.84 for HGDP (*p <* 0.001 for both). The slightly lower correlation observed in HGDP likely reflects increased STR genotyping noise due to differences in sequencing technology, as 1KGP was generated using newer high-coverage, PCR-free whole-genome sequencing (WGS) platforms that minimize amplification stutter artifacts and thereby improve STR genotyping accuracy. To provide an independent validation, we performed supervised population assignment using SNP- and STR-based genotypes from HGDP (Methods). The SNP-based model achieved near-perfect accuracy for continental population prediction (99%), while the STR-based model reached 91% accuracy when trained on raw genotypes (Fig. 3f). However, when STR genotypes were first reduced using PCA, the resulting accuracy became comparable to that of SNPs, demonstrating that STRs encode continental population structure consistent with SNPs and that appropriate feature transformation is also required to recover their full resolution when genotyping noise is elevated.

**Figure 3:**
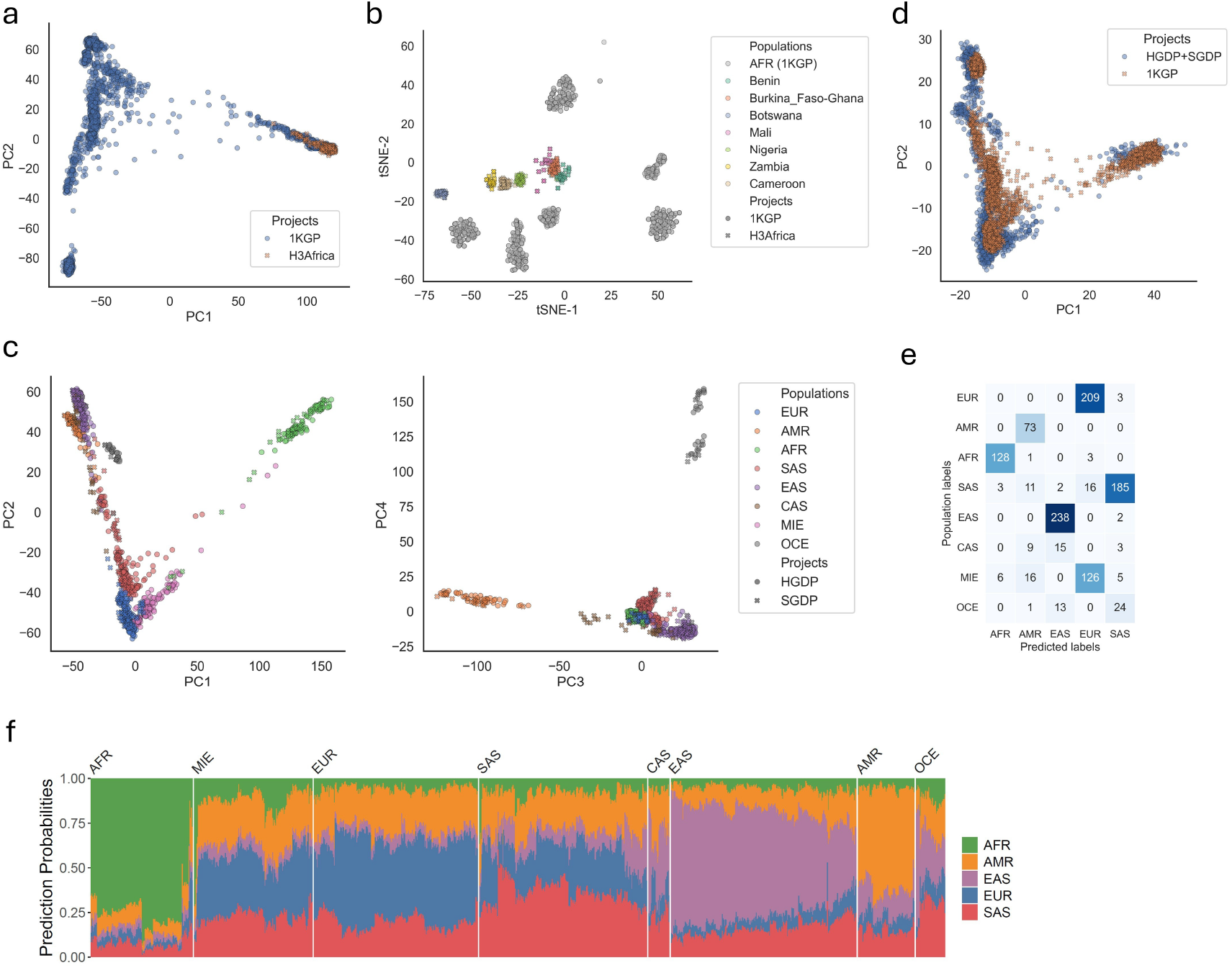
Cross-cohort validation and reproducibility of STR-based population structure. **a** Principal component analysis (PCA) of overlapping STR loci between the 1KGP and H3Africa datasets (PC1 vs. PC2) after batch effect correction. Each point represents an individual, colored and shaped by project. **b** t-SNE visualization of individuals based on the top 20 principal components from panel **a**. Only individuals from African populations are shown. Points are shaped by project. Individuals from African populations in the 1KGP are shown in gray, whereas those from the H3Africa dataset are colored according to seven regional population labels. **c** PCA of overlapping STR loci between the HGDP and SGDP datasets (PC1 vs.PC2 and PC3 vs PC4). Each point represents an individual, colored by population label and shaped by project. **d** PCA of overlapping STR loci between the 1KGP and HGDP+SGDP datasets (PC1 vs. PC2) after batch effect correction. Each point represents an individual, colored and shaped by project. **e** Confusion matrix of random forest–based continental population assignment trained on the 1KGP dataset and applied to the HGDP+SGDP dataset. The classification accuracy across shared continental populations (AFR, AMR, EAS, EUR, SAS) are 0.81. **f** Ancestry probabilities predicted by the random forest model trained on the 1KGP dataset and applied to the HGDP+SGDP dataset. Each column represents an individual, and stacked color segments indicate the predicted probabilities of assignment to each continental population from the 1KGP. Individuals are ordered by hierarchical clustering, and continental population labels from the HGDP+SGDP dataset are shown above for reference.

Taken together, these analyses demonstrate that STR- and SNP-based representations of population structure are concordant at the continental scale, whereas high-quality genome-wide STR variations uniquely enable high resolution of fine-scale regional population structure.

### STR-derived population structures are robust and reproducible across datasets

We next assessed the robustness and reproducibility of STR-derived population structure across independent datasets. The variable STR loci detected in each dataset were affected by differences in cohort composition, sequencing platforms, and genotyping pipelines. To enable cross-dataset comparisons, we first performed batch-effect correction on the shared STR loci (Methods). In the unsupervised setting, continental population structure was largely captured by the first two PCs. Therefore, after batch-effect correction, we first evaluated continental-level concordance by examining whether samples from corresponding continental groups overlapped in the first two PCs of the PCA constructed from these harmonized STRs. To further determine whether population structure was preserved across studies, we then used high-quality variable STR loci from the 1KGP dataset to train a random forest (RF) classifier for continental population assignment and validated its performance in three independent datasets: H3Africa, HGDP, and SGDP.

Since STR genotypes from 1KGP and H3Africa were generated using the same analytical pipeline, we first attempted to overlap the variable STR loci to assess cross-dataset consistency within African populations. As expected, over 90% of variable STRs were retained for joint analyses after batch effect correction. The complete overlap between the AFR population from the 1KGP and the H3Africa datasets confirmed that African population structures were reproducible between the two datasets (Fig. 3a; Supplementary Fig.S4a,b). Furthermore, the RF model trained on 1KGP accurately classified all H3Africa samples as African (AFR), confirming consistent ancestry representation across datasets. Using t-SNE based on the first 30 PCs, we further observed well-resolved regional population structure within African populations, showing clear separation between regional groups from 1KGP and H3Africa (Fig. 3b). This is consistent with the motivation for incorporating H3Africa, which was designed to address the underrepresentation of West, Central, and South African populations in the 1KGP dataset.

Next, we integrated STR genotypes from 1,092 samples in the combined HGDP and SGDP datasets, as more than 90% of variable STR loci were detected in both, likely due to the use of the same sequencing platform and genotyping pipeline. PCA of the shared STR loci demonstrated consistent continental population structure in the first four PCs, while dataset-associated separation was observed only in later PCs (Fig. 3c; Supplementary Fig.S4c). Hereafter we refer to this merged dataset as HGDP+SGDP (Methods) and evaluated its cross-dataset reproducibility relative to the 1KGP dataset. Intersecting STR loci across datasets yielded 22,933 overlapping loci, which exhibited noticeable differences (Supplementary Fig.S4d,e). For batch-effect correction, we focused on African (AFR), European (EUR), and East Asian (EAS) populations, which represent minimally admixed continental groups. Loci showing inconsistent average STR lengths among these three populations were removed (Supplementary Fig.S4d). The triangular arrangement of EAS, EUR, and AFR in PCA reflected the expected topology of human population divergence and provided a robust reference for evaluating cross-dataset consistency. After correction, 7,079 high-confidence loci were retained. PCA based on these harmonized loci showed clear overlap among continental groups (Fig. 3d; Supplementary Data 3), confirming that STR-derived population structures are robust across datasets generated from different sequencing and genotyping conditions. Additionally, we trained RF models using the harmonized STR loci from 1KGP for continental population assignment. The resulting confusion matrix showed high prediction accuracy (81%) for continental populations represented in both 1KGP and HGDP+SGDP datasets (Fig. 3e). Predicted probability profiles further illustrated meaningful population relationships: for example, Oceanian (OCE) samples, which were not present in the 1KGP, exhibited mixed probability profiles across multiple continental groups, reflecting their intermediate genetic affinities and mixed patterns of Oceanian population history (Fig. 3f; Supplementary Data 4).

These analyses demonstrate that STR-derived population structures are robust, reproducible, and transferable across diverse datasets. Despite differences in sequencing platforms, cohort compositions, and analytical pipelines, STR-based models consistently recapitulated the major continental population patterns observed in HGDP, SGDP, and H3Africa, each showing strong concordance with the reference population structure in 1KGP. The ability of 1KGP-trained models to generalize accurately to independent cohorts, and to infer plausible ancestry relationships for populations absent from the training set, underscores the stability of STR-based ancestry signals.

### Directional NMF decouples ancestry from STR mutational dynamics

Next, we sought to establish a framework to uncover the underlying structure of ancestral populations using genome-wide STRs. Based on the stepwise mutation model, STR allele lengths evolve through single-unit gains and losses with approximately equal probability. We therefore hypothesized that true ancestral population structures are encoded symmetrically in both the expansion and contraction directions of STR length mutations. To decouple ancestry signals from the intrinsic mutational properties of STRs, we developed Directional Non-negative Matrix Factorization (dNMF), a decomposition framework specifically tailored to infer ancestry coefficients from STR mutation dynamics. For each STR locus, repeat length was standardized across individuals using Z-score normalization. The resulting standardized matrix **D** ∈ R^*N*×*L*^, where *N* and *L* denote the number of individuals and STR loci, respectively, was then transformed into two directional components: **D**_*pos*_ = *max* (**D**, 0) in the expansion channel and **D**_*neg*_ = *max* (−**D**, 0) in the contraction channel. Two independent non-negative matrix factorizations were then fitted to each channel, yielding ancestry proportion matrices **W**_*pos*_ and **W**_*neg*_, and locus contribution matrices **H**_*pos*_ and **H**_*neg*_ in each channel. This formulation enables direct comparison of ancestry components derived from the expansion and contraction channels, allowing us to infer ancestral populations preserved across mutational directions and identify motif-specific biases associated with distinct mutation mechanisms.

We first applied dNMF to 74,465 variable STR loci from 3,202 samples in the 1KGP dataset and evaluated model stability across a range of component numbers (*K* = 3 to 25). The stability of the inferred ancestry components was assessed by averaging pairwise correlations of matched components across independent runs (Methods; Fig. 4a; Supplementary Fig.S5a,c,e). As the number of components (K) increased from 3 to 25, the number of shared components reached a maximum at *K* = 12 (Fig. 4b), where components across the expansion and contraction channels aligned in a one-to-one fashion. Beyond this point, the average correlation within each run declined sharply and became unstable, indicating that higher *K* values introduced component splitting and reduced internal consistency of the inferred ancestry structure. We therefore determined *K* = 12 as the optimal number of ancestry populations preserved across both mutation directions. At this optimal resolution, dNMF revealed ancestry components that recapitulated both major continental and regional population structures in the 1KGP dataset (Fig. 4g; Supplementary Data 5). Distinct components corresponded to the five major continental populations (AFR, AMR, EUR, SAS, EAS), while additional components captured substructure. Within AFR continental population, components *S*2, *S*6, *S*7, *S*8, *S*10, *S*12 differentiated West, East, and admixed African populations. European populations (EUR) were characterized by the components *S*1, *S*11, likely reflecting north-south gradients, whereas *S*3 and *S*9 captured analogous patterns within East Asian populations (EAS). Notably, the resolution of population structure inferred from STR-based exceeded that obtained from SNP-based admixture inference, which primarily resolved broad continental groupings. The STR-based dNMF model delineated subtle within-continent differentiation, particularly within the African population (AFR).

**Figure 4:**
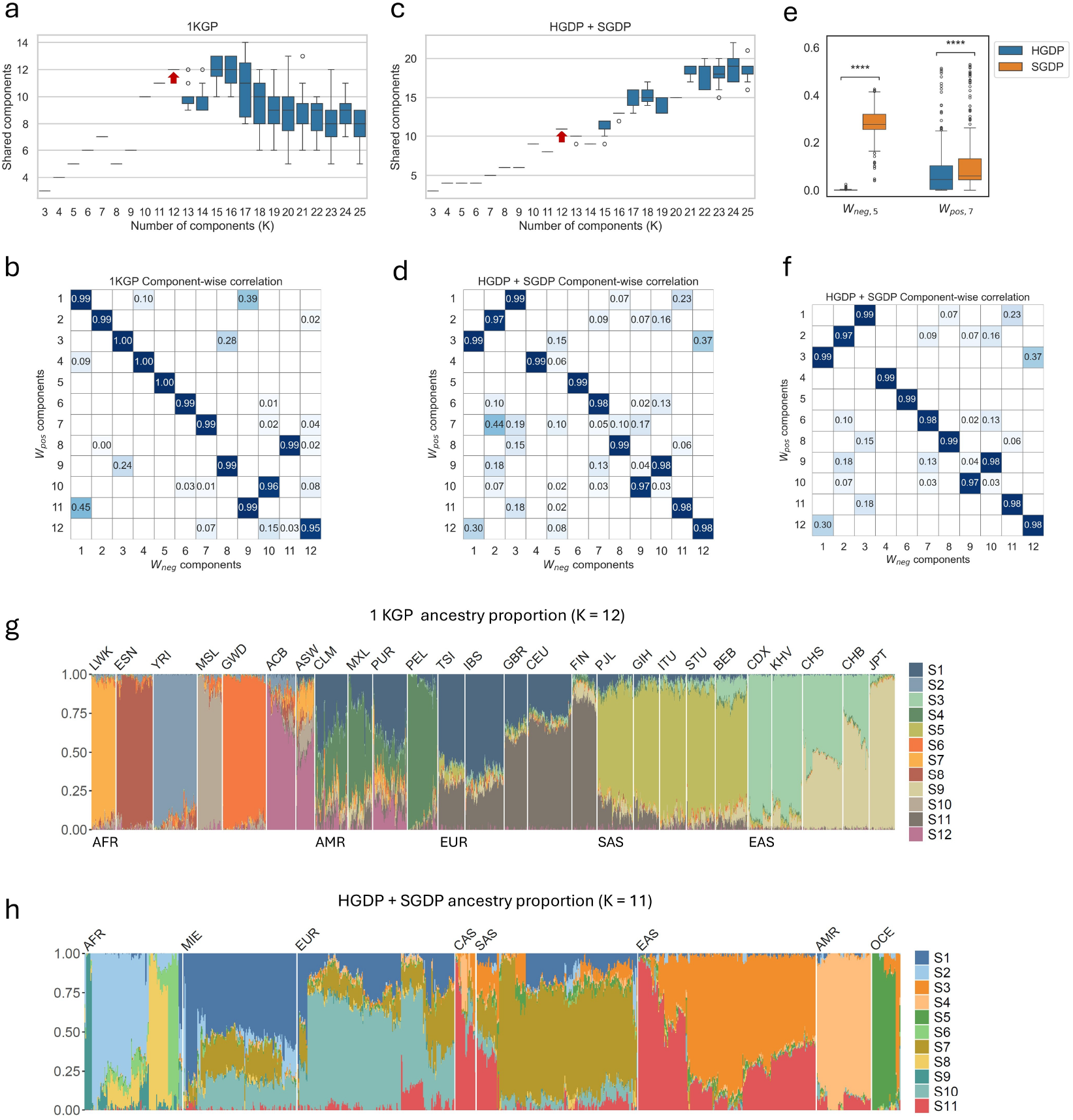
dNMF reveals robust and reproducible ancestry components across datasets. **a, c** Boxplots of the number of shared components across multiple runs in the 1KGP (a) and HGDP+SGDP (c) datasets. The x-axis indicates the number of components (*K*) used in the dNMF model. Each box shows the distribution of shared components identified across repeated runs for a given K. The central line indicates the median, the box boundaries represent the first and third quartiles, and whiskers extend to 1.5× the interquartile range. Outliers are shown as individual points. The optimal *K* is indicated by a red arrow (*K* = 12 for both datasets). **b, d** Heatmap showing pairwise Pearson correlation coefficients between normalized components (*K* = 12) from expansion (**W**_*pos*_) and contraction (**W**_*neg*_) channels in the 1KGP (b) and HGDP+SGDP (d) dataset. Each cell represents the Pearson correlation coefficient between component pairs, with color indicating the correlation strength. The corresponding correlation values are annotated in each cell. **e** Boxplots comparing normalized values of components **W**_*neg*,5_ and **W**_*pos*,7_ between the HDGP and SGDP datasets. Boxes indicate the interquartile range (IQR), horizontal lines represent the median, and whiskers extend to 1.5× the IQR. Outliers are shown as individual points. Statistical significance was assessed using the two-sided Mann–Whitney–Wilcoxon test (\*\*\*\**P <* 0.0001)^62^. **f** Heatmap showing pairwise Pearson correlation coefficients between normalized components (*K* = 11) in the HGDP+SGDP dataset after removing the components **W**_*neg*,5_ and **W**_*pos*,7_. **g, h** Normalized ancestry proportion using optimal *K* inferred by dNMF in the 1KGP (g, *K* = 12) and HGDP+SGDP (h, *K* = 11) datasets. Each column represents an individual, and stacked color segments indicate the proportion of each ancestry component derived from **W**_*pos*_. Individuals are ordered by hierarchical clustering. For the 1KGP dataset, continental population labels are shown below and regional population labels are shown above the plot. For the HGDP+SGDP dataset, continental population labels are shown above the plot.

Next, we applied dNMF to 63,746 variable STR loci of 1,092 samples from the HGDP+SGDP dataset. Using the same analytical procedure, we found that the average correlation among inferred components remained stable up to *K* = 12. In contrast to the 1KGP results, where the number of shared components started to decrease after reaching the maximum at *K* = 12, the results from HGDP+SGDP exhibited persistent but unstable component splitting at higher K values (Fig. 4c; Supplementary Fig.S5b,d,f), likely reflecting increased STR genotyping noise as discussed in the previous section. At *K* = 12, most ancestry components inferred from the expansion and contraction channels showed clear correspondence (Fig. 4d). However, two components lacked matching counterparts: **W**_*pos*,7_ and **W**_*neg*,5_. To examine whether these components were driven by technical artifacts, we calculated the Pearson correlation between their ancestry coefficients and the data source, dummy-coded as 0 (HGDP) and 1 (SGDP). We observed a very strong correlation for *W*_*neg*,5_ (*r* = 0.95, *P <* 10^−10^) and a weaker, though still significant correlation for *W*_*pos*,7_ (*r* = 0.22, *P <* 10^−10^) (Fig. 4e; Supplementary Fig.S6). These results indicate that *W*_*neg*,5_ was primarily driven by batch effects between datasets, whereas *W*_*pos*,7_ may be partially affected by dataset-specific artifacts. Both components were therefore excluded from subsequent cross-channel comparisons to ensure that only biologically meaningful ancestry signals were retained. After excluding the two components from each channel, the remaining ancestry components displayed strong directional concordance (Fig. 4f). We therefore determined *K* = 11 as the optimal number of ancestry populations in the HGDP+SGDP dataset (Fig. 4h; Supplementary Data 6). The ancestry proportions inferred from the HGDP+SGDP dataset closely resembled those identified in the 1KGP dataset. Two major ancestral populations were recovered within East Asian (EAS) and European populations (EUR), while four distinct ancestry components were observed within the Africa populations (AFR). Beyond continental patterns, 141 regional populations displayed unique ancestry profiles (Supplementary Fig.S7), demonstrating that dNMF effectively captures both broad continental patterns and regional population differentiation across globally diverse cohorts.

The separation of mutational directions allows the model to distinguish genuine biological signals from technical artifacts, as batch-associated components typically manifest asymmetrically across channels. Therefore, despite the elevated genotyping noise in the HGDP+SGDP dataset, the core ancestry components recovered at the optimal resolutions (*K* = 12 for 1KGP and *K* = 11 for HGDP+SGDP) remained highly concordant between the two datasets. These results confirm that STR-based dNMF accurately resolves the number of ancestral populations and reliably estimates population admixture structure.

### Ancestry-informative STR signatures and motif specialization

STR mutation rates and patterns are known to depend on repeat motif length and genomic context, suggesting that distinct motif classes and genomic distributions may differentially encode population-specific information. To investigate how individual STR loci contributed to the ancestry components inferred by the dNMF, we first examined the aligned locus contribution matrices **H**_*pos*_ and **H**_*neg*_. Across both the 1KGP and HGDP+SGDP datasets, we observed moderate negative correlations between aligned locus contributions from the two channels (*r* = −0.6 to −0.3; Fig. 5a,b, left), indicating that loci contributing strongly in one channel tended to have weak contributions in the other channel. This suggests that STR expansions and contractions encode consistent ancestry information through opposing mutational processes, further supporting the hypothesis that population structure is independent of the STR mutation direction.

**Figure 5:**
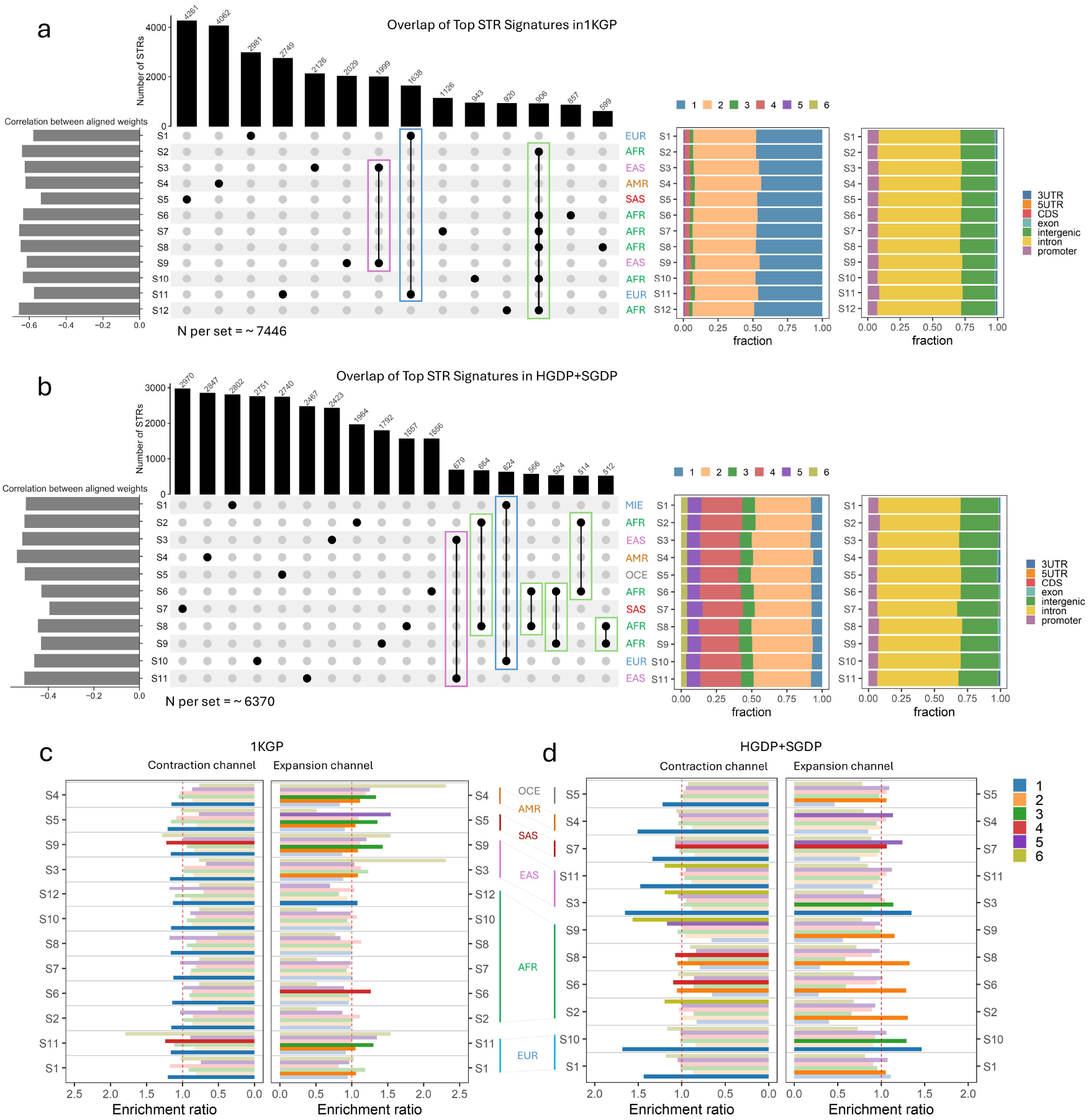
Ancestry-informative STR signatures and motif enrichment patterns revealed by dNMF. **a, b** Barplot in the left panel shows Pearson correlation coefficients between aligned weights (**H**_*pos*_ and **H**_*neg*_) for each ancestry component. Upset plot in the middle panel show overlap of the top 5% ancestry-informative STR signatures of aligned ancestry components from both channels. Bars above indicate the number of unique STRs per ancestry component, and connected dots represent shared loci between components. Categories with fewer than 500 STRs were excluded. Population labels denote the dominant population associated with each component. Right panels summarize motif length compositions and genomic annotation distributions of STR signatures. **c, d** Motif enrichment analysis of ancestry-informative STR signatures in the contraction (left) and expansion (right) channels for the 1KGP (c) and HGDP+SGDP (d) datasets. Significant motif classes (1-6 bp) are indicated in bright colors. Enrichment ratios were computed using Fisher’s exact test, with multiple testing correction performed using the Benjamini-Hochberg^63^ procedure (*FDR <* 0.05).

To further characterize the ancestry-informative STR signatures, we identified the top 5% of loci with the highest contributions within each ancestry component from aligned contribution matrices **H**_*pos*_ and **H**_*neg*_ (Supplementary Fig.S9,10; Supplementary Data 7,8). We assessed the specificity and overlap of these ancestry-informative STR signatures (Fig. 5a,b, middle). In both datasets, the majority of STR signatures were unique to individual ancestry components, indicating strong population specificity. Notably, overlapping STR signatures were primarily shared among closely related populations, such as the two ancestral components within Europe (EUR) and the two within East Asian (EAS), and several components within Africa (AFR). These overlaps likely reflect recent shared ancestry and genetic continuity. In contrast, components representing distinct continental populations (SAS) showed minimal sharing, consistent with long-term divergence and demographic isolation. Together, these results demonstrate that dNMF-derived STR signatures capture both the specificity and hierarchical relationships of human population structure at the locus level.

To explore the underlying mutational basis of these STR signatures, we compared their motif compositions and genomic distributions in both datasets (Fig. 5a,b, right). In the 1KGP dataset, most ancestry-informative STR signatures consisted of 1-bp and 2-bp motifs, whereas in the HGDP+SGDP dataset, 2-bp and 4-bp motifs were more prevalent. Quantitatively, 1-bp and 2-bp motifs accounted for 92.9% of variable STRs in the 1KGP, while 2-bp and 4-bp motifs comprised 69.7% of variable STRs in the HGDP+SGDP (Supplementary Fig.S8a,b). These differences largely reflect variation in the motif composition of the input STR sets rather than differences in populationinformative signals. Despite differences in motif frequency, the genomic distributions were highly consistent across components and datasets, with most STR signatures located in intronic and intergenic regions, consistent with their predominantly neutral evolutionary processes. Additionally, we performed motif enrichment analysis on directionally biased STR signatures identified in each ancestry component from the expansion and contraction channels. Motif enrichment ratios were computed for the top 5% of loci contributing to each component, and statistical significance was assessed using Fisher’s exact test with FDR correction (*FDR <* 0.05). Significant enrichments are highlighted as bright-colored bars (Fig. 5c,d). Notably, homopolymeric repeats were significantly enriched across all ancestry components from the contraction channel in the 1KGP and in more than half of the components from the contraction channel in the HGDP+SGDP dataset, whereas dinucleotide repeats were enriched in nearly half of the components from the expansion channel in both datasets (Supplementary Fig.S11,12). These directional biases likely arise from genomewide mutational tendencies rather than locus-specific selective pressures. Consistent with this, no significant enrichment was detected for STR signatures in the genic or regulatory regions (Supplementary Fig.S8c,d), showing that ancestry-informative STRs are broadly distributed across the genome and reflect primarily neutral, mutation–drift–driven processes.

Finally, we evaluated whether STRs of different motif lengths could independently uncover population admixture. To test this, we partitioned STRs by motif length (1–5 bp) in the 1KGP, applied dNMF to each subset separately, and determined the optimal number of ancestry components *K*. The resulting numbers of inferred ancestral populations were 12, 10, 6, 6, and 5 for 1-bp to 5-bp STRs, respectively (Fig. 6; Supplementary Fig.S13). Despite differences in motif length, the inferred continental and regional population patterns were broadly concordant across motif classes, indicating that population structure is consistently encoded across multiple mutational scales. Notably, shorter motifs (1–2 bp) captured finer-scale substructure, particularly within African populations, whereas longer motifs (3–5 bp) delineated broader continental groupings, reflecting deeper evolutionary divergence. These findings suggest that distinct motif classes encode complementary layers of population history, corresponding to the hierarchical nature of human demographic evolution.

**Figure 6:**
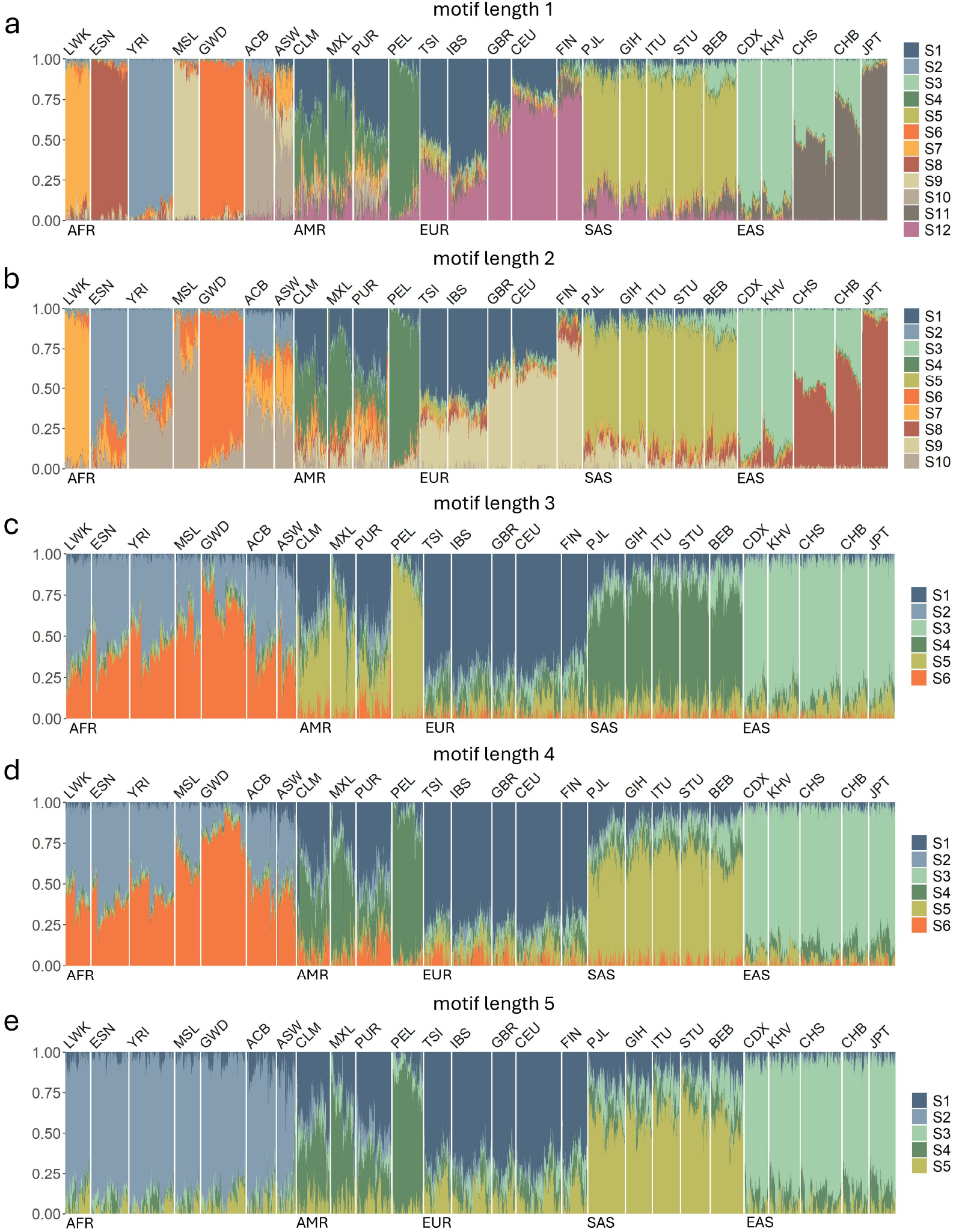
Motif-specific admixture inference reveals hierarchical encoding of population structure. **a–e** Normalized ancestry proportions inferred by dNMF using STRs stratified by motif length (1–5 bp) in the 1000 Genomes Project (1KGP). For each STR motif subset, dNMF was applied independently to estimate ancestry coefficients at the optimal number of ancestry components (*K* = 12, 10, 6, 6, and 5, respectively). Individuals are ordered by hierarchical clustering. Continental population labels are shown below and regional population labels are shown above the plot.

## Discussion

This study establishes STRs as powerful genome-wide markers for inferring human population structure. Although STRs have often been regarded as too mutable or technically challenging for population-level analyses, our results demonstrate that genome-wide STR variation encodes stable and interpretable signals of human ancestry across both continental and regional scales. We introduce a comprehensive STR-based framework that integrates unsupervised clustering, supervised classification, and a novel population admixture inference model, Directional Non-negative Matrix Factorization (dNMF). Through this multi-model approach, we show that STR variation reveals robust and reproducible patterns of population structure across multiple large-scale human genomic datasets. Beyond accurately estimating fine-scale latent ancestry components, dNMF also uncovers distinct mutational mechanisms underlying STR evolution. Together, these findings position STRs as a complementary and biologically informative class of population markers capable of capturing fine-scale demographic differentiation while reflecting the mutational processes that shape human genetic diversity.

Using genome-wide variable STRs derived from whole-genome sequencing, our study demonstrates that STRs offer distinct advantages over SNPs for population structure inference, revealing patterns that may be undetectable using biallelic markers. Previous comparisons of population structure inferred from STRs and SNPs were limited to hundreds of STR loci contrasted against millions of SNPs^38,39^, thereby missing the full spectrum of genome-wide variation captured in this study. Here, the strong concordance between STR- and SNP-based continental population patterns across datasets confirms that both variant types capture the same global population structure, while the finer differentiation observed in STR-based analyses underscores their sensitivity to recent lineage diversification. Notably, the accuracy of the STR-based model in supervised population assignment reached 99% at the regional level in the 1KGP dataset, substantially exceeding the 82% accuracy achieved by the SNP-based model. A separate study using 4.5 million SNPs reported a mean prediction accuracy at 88.61% (80% training and 20% testing) for 26 regional populations in the 1KGP dataset^40^, further highlighting the high efficiency and resolution of STR-based models. Beyond their intrinsic informativeness, STR-based models also generalize across datasets and sequencing platforms. Recent efforts have been made to harmonize the deep sequenced human genome from the 1KGP and HGDP^41^, focusing on identifying high-quality SNV, indels and structure variants. Similarly, we showed that after batch-effect correction, random forest classifiers trained on 1KGP data accurately predicted continental populations in the H3Africa and HGDP+SGDP datasets. This cross-dataset consistency indicates that STR-based ancestry models can be transferred between studies in a manner analogous to SNP-based pipelines, enabling scalable ancestry inference in diverse human populations. The harmonized STR loci identified in this study may thus serve as a reference set for future comparative human population analyses based on STR variations.

Building on the stepwise mutation model of STR evolution^37,42^, we developed a novel framework, dNMF, which explicitly exploits the mutation dynamics of STRs to estimate ancestry coefficients. While SNP-based methods typically determine the optimal number of *K* components by minimizing cross-validation error, dNMF instead enables the estimation under the hypothesis that true and stable ancestral populations are encoded consistently across bidirectional STR mutations. Applied to the 1KGP and HGDP+SGDP datasets, dNMF determined the optimal ancestry components at *K* = 12 and *K* = 11, respectively. The inferred ancestry patterns were highly concordant between datasets, indicating stable and reproducible population structure. Notably, these values exceed the typical numbers inferred from SNP-based methods, which generally reflect major continental populations. For instance, analyses of the harmonized 1KGP and HGDP datasets have reported a best-fit value of *K* = 6 using ADMIXTURE^41^, while the SGDP dataset was reported to have an optimal value of *K* = 5 using ADMIXTURE^35^. More recent scalable admixture models such as TetraStructure^4^ and SCOPE^43^ selected *K* = 10 latent populations for the HGDP dataset; however, the resulting ancestry patterns were noisier and less biologically interpretable compared with those from dNMF. Furthermore, when applied to the combined HGDP+SGDP dataset, dNMF successfully detected and distinguished technical artifacts arising from differences in sequencing protocols, demonstrating its robustness to moderate technical variation and its capacity to disentangle true population structure signals from systematic bias. By accommodating the neutral and bidirectional nature of STR mutations, dNMF extends traditional SNP-based models to mutation-driven variation, providing a biologically interpretable representation of ancestry based on STR mutational dynamics. Recent work by Anya et al. has shown that simple DNA repeats evolve under a reproducible, mutation-driven equilibrium in which length-dependent expansion and contraction mutational biases maintain a stable repeat-length distribution across mammals^44^. These findings provide strong empirical support for the theoretical foundation of our framework.

In addition to the ancestry coefficient estimation, analysis of locus contribution matrices from the dNMF identified ancestry-informative STR signatures and revealed how different STR mutation mechanisms shape population structure. Despite the different composition of input STRs in the 1KGP and HGDP+SGDP datasets (Supplementary Fig.S8a,b), dNMF revealed robust and reproducible motif-specific enrichment patterns. The enrichment of homopolymers in the contraction channel and dinucleotide repeats in the expansion channel mirrors well-established mutational asymmetries reported in previous studies, where homopolymers are prone to contraction due to replication slippage and polymerase stalling^45,46^, while dinucleotide repeats, particularly AC motifs, frequently undergo expansion driven by strand misalignment and local DNA structural features such as Z-DNA formation^47–50^. The observation that these enrichments persist across diverse populations and datasets indicates that they arise from intrinsic, motif-dependent mutational dynamics rather than population-specific demographic history. Collectively, these results demonstrate that dNMF not only captures population structure but also systematically reflects the directional mutational tendencies of major STR motif classes at a genome-wide scale. Importantly, these enrichments were consistent across datasets and not preferentially localized to functional genomic regions, suggesting that ancestry informative STR variation largely reflects neutral mutation–drift processes. Furthermore, stratification by motif length revealed hierarchical ancestry encoding: short motifs (1–2 bp) captured fine-scale differentiation, particularly within African populations, while longer motifs (3–5 bp) delineated deeper continental divergences. This pattern implies that different STR classes record distinct temporal layers of human demographic history.

While this study demonstrates the power of genome-wide STRs for high-resolution population structure inference, several limitations remain. The accuracy and completeness of STR-based analyses depend strongly on sequencing coverage, read length, and genotyping precision. Current datasets, including the 1KGP, represent the highest-quality STR resources available from short-read sequencing. However, further advances in long-read sequencing and genotyping algorithms will be essential to extend STR-based population analyses to more diverse cohorts and to structurally complex regions of the genome. As sequencing technologies and genotyping algorithms continue to improve^51^, integrating high-quality STRs with SNPs could enable multi-layered reconstructions of population history across time scales. Moreover, extending dNMF to model local ancestry or haplotype structure may allow joint inference of mutation directionality and recombination, yielding finer resolution of evolutionary processes. Beyond human populations, these principles can be generalized to other species^44,52–54^, establishing STRs as broadly applicable markers to understand how mutational processes shape genetic diversity across evolution.

Compared with SNPs, STRs provide a richer mutational alphabet that encodes both the magnitude and direction of evolutionary change, enabling new ways to model population structure beyond binary genotype data. The dNMF framework introduces a conceptually distinct approach to ancestry inference by treating mutation directionality as an informative dimension rather than noise. The methodological and conceptual advances presented here, particularly dNMF, open new avenues for integrating mutational dynamics into demographic inference and for connecting molecular mutation processes with the broader narrative of human evolutionary history.

## Methods

### Dataset collection

STR genotype data in Variant Call Format (VCF) for 3,202 samples from the 1000 Genomes Project (1KGP) and 348 samples from the H3Africa Project were obtained from version II of the EnsembleTR calls (https://github.com/gymrek-lab/EnsembleTR)^24^ . Autosomal STR loci genotyped using HipSTR^25^ and with motif lengths of 1–6 base pairs were retained for downstream analyses.

High-coverage whole-genome sequencing (WGS) data in CRAM format for 828 samples from the Human Genome Diversity Project (HGDP) and 276 samples from the Simons Genome Diversity Project (SGDP) were obtained from the International Genome Sample Resource (IGSR)^55^. These datasets were then used for STR genotyping.

SNP genotype data for 1KGP were retrieved from the official 1KGP high-coverage phased release (https://ftp.1000genomes.ebi.ac.uk/vol1/ftp/data_collections/1000G_2504_high_coverage/working/20220422_3202_phased_SNV_INDEL_SV/). SNP genotype data for HGDP were downloaded from the Sanger Institute release (https://ngs.sanger.ac.uk/production/hgdp/hgdp_wgs.20190516/). SNPs were filtered using PLINK^56^ with parameters `--geno 0.1 --maf 0.1 --indep-pairwise 50 10 0.1` to remove variants with low call rate, low minor allele frequency, and high linkage disequilibrium. After filtering, 144,912 polymorphic SNPs were retained for 1KGP and 127,384 for HGDP.

### STR genotyping and quality control

For the HGDP and SGDP datasets, STR genotyping was performed for each sample using HipSTR v0.6.2 with the following non-default parameters: `--min-reads 5 --max-reads 2000000`, us-ing the hg38 STR reference available at https://github.com/HipSTR-Tool/HipSTR-references/blob/master/human/hg38.hipstr_reference.bed.gz. Sample-level VCF files were then filtered using dumpSTR^57^ v3.0.3 with the following options: `--min-locus-callrate 0.80 --min-locus-hwep 0.000001 --filter-regions hg38 segdup.sorted.bed.gz --filter-regions-names SEGDUP`. Segmental duplication regions were obtained from the UCSC Genome Browser^58^. Filtered sample-level VCF files were merged within each dataset using a modified version of mergeSTR^57^ (https://github.com/gymreklab/TRTools/tree/conf_ref) and subsequently corrected using the Python script ((https://github.com/gymreklab/1000Genomes-STRs/blob/main/Hipstr_correction.py). For each dataset, STR loci called in fewer than 75% of samples were excluded. Only autosomal loci were retained for analysis. For each retained locus, the mean allele length was calculated per individual as the average of the two alleles reported by HipSTR. Variable STR loci were further filtered using different length variation thresholds (Supplementary Data 1).

### Unsupervised clustering to genotype data

For comparison, we constructed an STR genotype matrix **D**_*m,n*_ and an SNP genotype matrix **P**_*s,n*_, where *m* and *s* represent the number of variable STRs and polymorphic SNPs respectively, and *n* denotes the number of samples. Each entry in **D**_*m,n*_ corresponds to the standardized mean allele length of a given STR locus in a specific individual, whereas each entry in **P**_*s,n*_ represents the biallelic genotype dosage (0, 1, or 2) for a given SNP. Missing STR genotypes were set to NaN and then imputed using the locus-specific mean. For the 1KGP dataset we used *m* = 31,359 variable STR loci (length variation *>* 5) and *s* = 144,912 polymorphic SNPs for 3202 samples and performed principal component analysis (PCA) on both variant types. To evaluate continental and regional population structure, we applied k-means clustering to the first N principal components (2–30 PCs), where N varied across experiments. The number of clusters was set to match the known numbers of continental or regional populations. Clustering performance was assessed using the Adjusted Rand Index (ARI) against the corresponding population labels. To visualize regional population substructure, we applied the t-distributed stochastic neighbor embedding (t-SNE) algorithm to the top 30 principal components derived from STR-based and SNP-based PCA. For the HGDP dataset, we used a subset of samples (*n* = 802) with both STR and SNP genotypes available, consisting of *m* = 34,911 variable STR loci (length variation *>* 5) and *s* = 127,384 polymorphic SNPs. PCA was then performed in the same way on each variant type. All clustering methods were implemented using the Python scikit-learn library^59^ v1.6.1.

### Supervised population assignment

To evaluate the predictive power of STRs for population assignment, we trained random forest classifiers (RF) separately on STR and SNP genotype data from the 1000 Genomes Project (1KGP). The same genotype matrices used in the unsupervised clustering analysis served as input for these supervised models. For STRs, models were trained directly on the imputed and standardized STR genotype matrix without dimensionality reduction. To evaluate potential confounding effects of related individuals, models were trained and tested both with and without the 698 related samples documented in the 1KGP metadata. Classification accuracies were compared in these two settings to assess the robustness of STR-based model to familial structure. Since model performance remained unchanged when related individuals were excluded, we concluded that familial structure did not confound the analyses and therefore retained all samples for subsequent comparisons. For SNPs, models were trained using the top 30, 40, and 50 principal components (PCs) derived from the SNP genotype matrix, which capture the major axes of genetic differentiation while reducing the high dimensionality and sparsity of raw SNP genotypes. Model performance was evaluated using stratified five-fold cross-validation at both continental and regional population levels. Classification accuracies were computed for each fold, and mean accuracies were reported across all folds. To further assess the robustness of the STR-based model, we additionally trained multinomial naïve Bayes models (`GaussianNB`).

To evaluate the robustness of STR-based models and genomic distribution of informative STRs, we assessed classification accuracy as a function of (i) the number of downsampled STR loci and subsets of variable STRs available on each chromosome, using the set of 31,359 variable STRs from the 1KGP dataset:

- **Downsampling of STR loci**. This analysis was designed to assess the minimum number of STR markers required to recover continental and regional population structure. We randomly sampled subsets containing 5000, 4000, 3000, 2000, 1000, 500, 100, and 50 loci. For each target size, 10 independent downsampling replicates were generated with replacement. An RF classifier was trained for each replicate, and classification accuracy was evaluated using stratified five-fold cross-validation at both continental and regional population levels. Accuracies across replicates were recorded for each marker set size.
- **Chromosome-specific STR subsets**. To examine the genomic distribution of ancestryinformative STRs, we constructed chromosome-specific STR genotype matrices for each autosome using all variable STR loci present on that chromosome. The number of variable STRs per chromosome ranged from 400 (chromosome 21) to 2,650 (chromosome 2), with an average of 1,425 loci across autosomes. For each chromosome-specific STR genotype matrix, RF classifiers were trained and evaluated using stratified five-fold cross-validation at the continental and regional levels.

All RF classifiers were trained with 100 trees (`n estimators=100`), a maximum tree depth of 50 (`max depth=50`), a minimum of 10 samples required to split an internal node (`min samples split=10`), and a minimum of 5 samples at each leaf node (`min samples leaf=5`). All other parameters were set at their default values. Hyperparameters were selected based on preliminary tuning to balance performance and computational efficiency. All models were implemented using the Python scikit-learn library^59^ v1.6.1.

### Genetic distance calculation and hierarchical clustering

We measured genetic differentiation using STR-based and SNP-based pairwise genetic distances among individuals from the 1KGP and HGDP datasets. For both datasets, STR and SNP genotype data were the same sets as those used in the unsupervised clustering analyses. STR-based pairwise genetic distances were calculated using the Goldstein distance^37^, which quantifies repeatlength differences under the stepwise mutation model. SNP-based distances were calculated as one minus the proportion of shared alleles between individuals using PLINK. Hierarchical clustering of each distance matrix was performed using the Ward linkage method implemented in the `scipy.cluster.hierarchy` module (v1.12.0). To evaluate the overall similarity between STR- and SNP-based population structures, we conducted Mantel tests with 10,000 permutations to estimate empirical *P*-values. Pearson correlation coefficients (r) between the two distance matrices were used as the test statistic.

We then obtained the geographic coordinates for each regional population in both datasets. Geographical distances between regional populations were computed using the Haversine formula, assuming an Earth radius of 6,371 km. Regional-level genetic distances were calculated by averaging all pairwise individual level distances across each population pair. Finally, we compared the correlations between geographical distances and STR-based or SNP-based genetic distances within each continental population to evaluate how well each variant type reflects spatial patterns of human population structure.

### Overlapping STR identification between datasets and batch-effect correction

To maximize the number of comparable loci across datasets, the set of variable STRs in the 1KGP was first expanded to 74,465 loci by including all loci exhibiting length variation greater than 2. Among these, approximately 95% were also detected in the filtered STR genotype dataset from H3Africa, indicating substantial overlap and consistency in STRs between the two datasets. To assess and correct for potential batch effects, we calculated the absolute difference in average length between AFR samples in the 1KGP and H3Africa datasets. STR loci showing an absolute mean length difference greater than 1 were considered affected by systematic bias and were excluded from the subsequent analyses. Overall, 68,579 STR loci were retained for the integrated 1KGP and H3Africa datasets.

The HGDP dataset included 828 samples, and the SGDP dataset included 276 samples. We identified 12 duplicate individuals present in both datasets; for these, the HGDP entries were retained to ensure consistency of metadata and genotyping format. The two datasets were then merged into a combined reference panel, referred to as HGDP+SGDP. For continental population labels, populations in the HGDP were labeled as EAS, SAS, MIE, EUR, AFR, AMR, and OCE, corresponding to East Asia, Central South Asia, Middle East, Europe, Africa, America, and Oceania, respectively. Populations in the SGDP were labeled as AFR, AMR, EAS, OCE, EUR, CAS, and SAS, representing Africa, America, East Asia, Oceania, West Eurasia, Central Asia Siberia, and South Asia, respectively. Overlapping variable STR loci between HGDP and SGDP were identified using the pyranges^60^ module (v0.1.4) based on matching chromosome coordinates and STR start and end positions. In the resulting HGDP+SGDP dataset, 63,746 STR loci from 1,092 individuals were retained for downstream analyses.

We further identified overlapping STR loci between the 1000 Genomes Project (1KGP) and HGDP+SGDP datasets, yielding a total of 22,933 shared loci. 23 individuals were present in both datasets, and we excluded them from the 1KGP dataset. Joint PCA of these loci revealed substantial technical artifacts indicative of batch effects between datasets. To correct for these discrepancies, we calculated the absolute differences in mean allele length between AFR, EUR, and EAS samples from the two datasets. These three populations were selected because they represent relatively unadmixed continental groups, providing a stable reference for cross-cohort comparison. STR loci showing an absolute mean length difference greater than 1 were considered inconsistent and excluded from further analyses. After this correction, 7,079 STR loci were retained, and the major continental populations showed clear overlap in the joint PCA, indicating effective batch-effect removal and alignment of continental population structure between datasets.

### Cross-dataset ancestry prediction

To evaluate the reproducibility of STR-derived population structures across datasets, we trained independent RF classifiers on the 1KGP dataset and applied them to predict continental population labels in the H3Africa and HGDP+SGDP datasets. Each model was trained using the harmonized STR loci identified as overlapping across datasets after batch-effect correction (see above). Model accuracy was first estimated by five-fold cross-validation within the 1KGP dataset prior to crosscohort prediction. The trained model was then applied to the H3Africa and HGDP+SGDP samples to predict population labels and estimate ancestry probabilities. The RF model was implemented using the same parameters and configuration as described previously. Prediction performance was evaluated using confusion matrices and per-class probability distributions to assess the degree of concordance between predicted and true population labels. Populations not present in the 1KGP reference (e.g., OCE) were included to examine the model’s behavior under extrapolation. For these populations, predicted ancestry probabilities were compared across reference populations to determine whether unrepresented ancestries were assigned mixed or intermediate probability profiles, reflecting their relative genetic proximity to existing continental groups.

### Directional Non-negative Matrix Factorization (dNMF)

Directional Non-negative Matrix Factorization (dNMF) is a decomposition framework developed specifically to model the directional mutation dynamics inherent to STR evolution. The model extends the standard admixture framework^9^ to STR genotype data. It is based on the hypothesis that true ancestral population structures are robustly captured in both STR mutational directions, allowing for the identification of stable ancestral components and directionally biased STR loci.

### Input data preparation

The input to dNMF is a standardized STR genotype matrix **D** ∈ ℝ^*N*×*L*^, where *N* is the number of individuals and *L* is the number of variable STR loci. Each entry in **D** represents the mean allele length of an STR locus for an individual, normalized by the locus-specific mean and standard deviation. To separate STR variation by mutation direction, the standardized matrix was decomposed into two non-negative matrices:

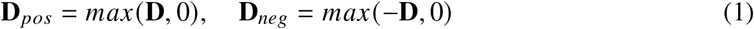

Here, **D**_*pos*_ contains positive *z*-scores corresponding to expansion-biased variation, while **D**_*neg*_ contains the absolute values of negative *z*-scores corresponding to contraction-biased variation.

### Model formulation

The expansion channel takes **D**_*pos*_ as input and contraction channel take **D**_*neg*_ as input. Each channel was independently factorized using the Non-negative Matrix Factorization (NMF), which approximates the input matrix as the product of two non-negative matrices by minimizing the Frobenius norm:

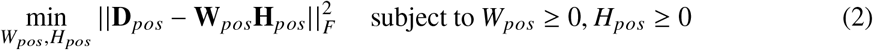

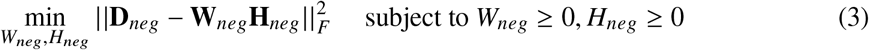

Optimization was performed using the coordinate descent solver, initialized with the non-negative double singular value decomposition (NNDSVD) to accelerate convergence and improve reproducibility. The resulting factorization can be expressed as:

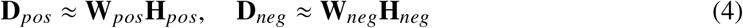

where **W**_*pos*_ and **W**_*neg*_ are *N* × *K* component matrices, and **H**_*pos*_ and **H**_*neg*_ are *K* × *L* locus contribution matrices, with *K* denoting the number of components. Each component matrix was then row-normalized to represent ancestry proportion matrices:

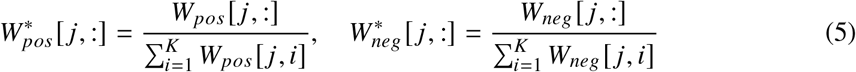

### Component alignment and stability assessment

To align ancestry components between channels, a *K* × *K* correlation matrix **C** was constructed, where the cell *c*_*i j*_ is the Pearson correlation coefficient between the *i*-th column vector of 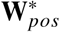 ancestry component *i* from the expansion channel) and the *j* -th column vector of 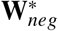 (ancestry component *j* from the contraction channel). The Hungarian algorithm was applied to the correlation matrix **C** to find the optimal one-to-one assignment 𝒫 that maximizes the average correlation:

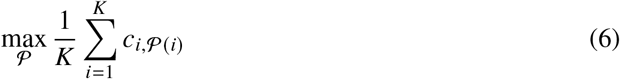

A matched pair (*i*, 𝒫(*i*)) was considered as shared and stable ancestry component if the corresponding correlation coefficient *r*_*i*,𝒫(*i*)_ satisfied the stringent criterion:

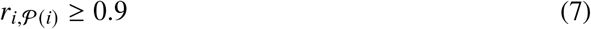

Average correlations and the number of shared components were recorded for each value of *K* across ten independent runs. Reconstruction errors from each channel were also monitored for comparison (Supplementary Fig.S5). The optimal number of ancestry components *K*_opt_ was determined as the value at which the number of shared components reached its maximum before the average correlation became unstable and began to decline. All procedures described above were applied independently to the 1KGP and HGDP+SGDP datasets to assess the stability, reproducibility, and transferability of the inferred ancestry components.

### Identification and characterization of ancestry-informative STR signatures

Ancestry-informative STR signatures were derived from the locus contribution matrices **H**_*pos*_ and **H**_*neg*_ obtained from the dNMF decomposition at the optimal number of components *K*_*opt*_. The two matrices were aligned using the optimal mapping 𝒫 determined by the Hungarian algorithm. Pearson correlation coefficients were then calculated between matched loci across the expansion and contraction channels to assess directional concordance. For each ancestry component, STR loci were ranked by their contribution values, and the top 5% were defined as ancestry-informative STR signatures. STR signatures from the contraction channel corresponding to aligned ancestry components were merged with those from the expansion channel. In total, approximately 10% of all input STR loci were retained as ancestry-informative STR signatures for each ancestry component.

To quantify the specificity and overlap of STR signatures among ancestry components, intersections were computed across all component-specific signature sets, and the results were visualized using UpSet plots. Motif composition and genomic distributions of STR signatures were characterized by intersecting STR genomic coordinates with gene annotations from the GENCODE v48 database using Bedtools^61^ v2.31.1. STRs were categorized by repeat motif length (1–6 bp) and genomic location (3’UTR, 5’UTR, CDS, non-coding exon, promoter[TSS +/-1kb], intergenic, intron). For each dataset, the relative proportion used all input STRs as the background distribution. Directional bias was assessed by comparing the expansion- and contraction-biased signature sets from **H**_*pos*_ and **H**_*neg*_. Enrichment analysis was performed using Fisher’s exact test with FDR correction to identify overrepresented motif classes or genomic categories (two-sided test, *FDR <* 0.05).

### Evaluation of ancestry resolution across motif classes

To evaluate the ancestry resolution of different STR motif classes, STR loci from the 1KGP dataset were used. The 1KGP STR genotype dataset was selected for this analysis due to its high sequencing quality and stringent quality control, which provide reliable genotyping accuracy across motif types. STR loci were stratified by repeat motif length (1–6 bp), resulting in 36,111, 33,032, 2,690, 1,724, 830, and 78 loci for 1-bp through 6-bp STRs, respectively. The dNMF model was then applied independently to each motif subset, using the same preprocessing and normalization procedures as described above. For each motif class, the number of ancestry components *K* was optimized based on model stability, assessed by the average correlation of ancestry components across ten independent runs. The inferred optimal number of ancestry components *K*_*opt*_ for each motif class was recorded to assess how motif length influences the hierarchical resolution of population structure.

## Supporting information

Description of supplementry data

## Acknowledgements

The computational work in this study was performed on the Earth cluster operated by the HPC team at the School of Life Sciences and Facility Management, Zürich University of Applied Sciences. The authors thank Dr. Hangjia Zhao for valuable input and constructive discussions.

## Author contributions

F.X. and M.A. conceived the project. F.X. curated the datasets, performed the data analysis, developed the methods, and generated the visualizations. M.A. and M.M. supervised the research. F.X. wrote the initial manuscript draft. All authors discussed the results and contributed to the revision of the final manuscript.

## Data availability

STR genotypes for the 1000 Genomes Project (1KGP) and H3Africa samples used in this study are available at the EnsembleTR GitHub repository [https://github.com/gymrek-lab/EnsembleTR]. STR genotypes generated from the Human Genome Diversity Project (HGDP) and Simon Genome Diversity Project (SGDP) in this study are available on GitHub repository [https://github.com/acg-team/STR_population_structure]. The WGS datasets for the HGDP and SGDP samples used in this study are available from the International Genome Sample Resource (IGSR) (https://www.internationalgenome.org/data-portal/data-collection). All analyses are based on the GRCh38 reference genome [https://storage.googleapis.com/genomics-public-data/resources/broad/hg38/v0/Homo_sapiens_assembly38.fasta]

## Code availability

Code used in this study is available on GitHub (https://github.com/acg-team/STR_population_structure).

## Supplementary materials

Supplementary Figures S1 to S13

Supplementary Data 1-8

