## Supplementary material for "High-resolution population structure inference using genome-wide short tandem repeat variations": Description of supplementry data

### Description of Additional Supplementary Files

File Name: Supplementary Data 1

Description: Numbers of variable STR loci within and across datasets

File Name: Supplementary Data 2

Description: Sample information for the HGDP+SGDP dataset

File Name: Supplementary Data 3

Description: Harmonized STR loci between the 1KGP and HGDP+SGDP datasets

File Name: Supplementary Data 4

Description:  Predicted ancestry probabilities for the HGDP+SGDP dataset

File Name: Supplementary Data 5

Description: Normalized Ancestry proportions inferred by dNMF for the 1KGP dataset

File Name: Supplementary Data 6

Description: Normalized Ancestry proportions inferred by dNMF for the HGDP+SGDP dataset

File Name: Supplementary Data 7

Description: Ancestry-informative STR signatures per component in the 1KGP

File Name: Supplementary Data 8

Description: Ancestry-informative STR signatures per component in the HGDP+SGDP
