## Supplementry figures for "High-resolution population structure inference using genome-wide short tandem repeat variations"

### This PDF file includes:

Supplementary Figures S1 to S13

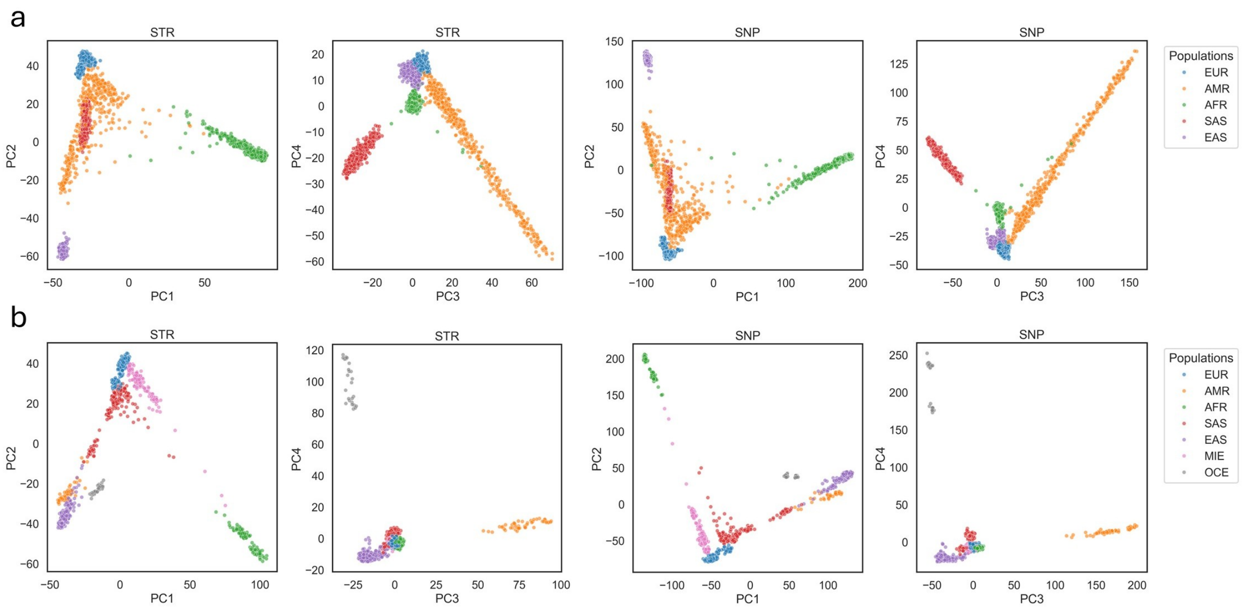


**Figure S1**: **Principal components analysis using variable STRs and polymorphic SNPs.** **a**. PCA was performed using STRs and SNPs in the 1KGP. The projections of each sample along PC1 and PC2, PC3 and PC4 are shown. **b**. PCA was performed using STRs and SNPs in the HGDP. The projections of each sample along PC1 and PC2, PC3 and PC4 are shown. Each point represents a single individual. Points are colored based on the continental population labels.

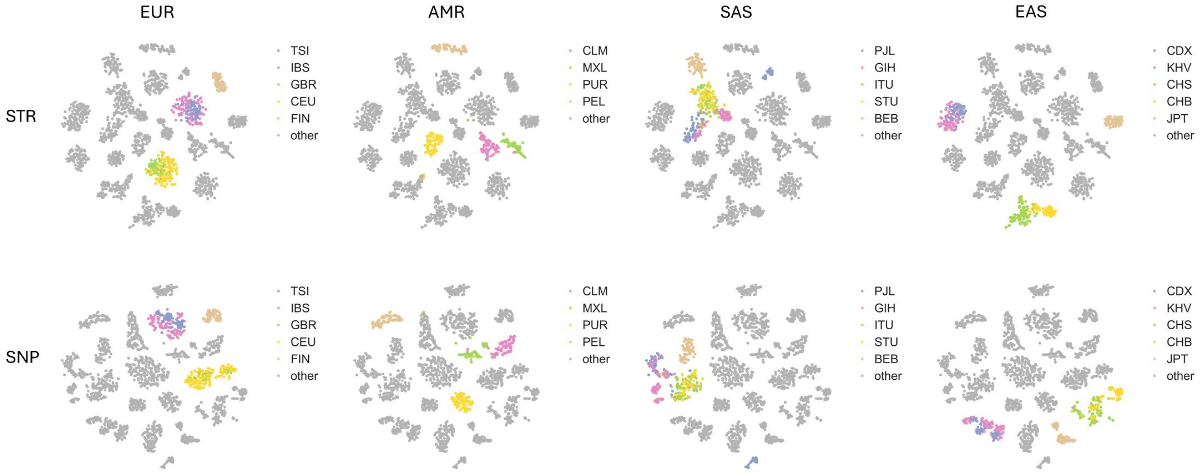


**Figure S2**: **t-SNE visualization of non-African samples using the top 30 principal components (PCs).** PCs were derived from STR-based (upper panel) and SNP-based (lower panel) PCA results. For each continental population, samples belonging to the focal population are colored by regional population labels, while all other samples are shown in grey.

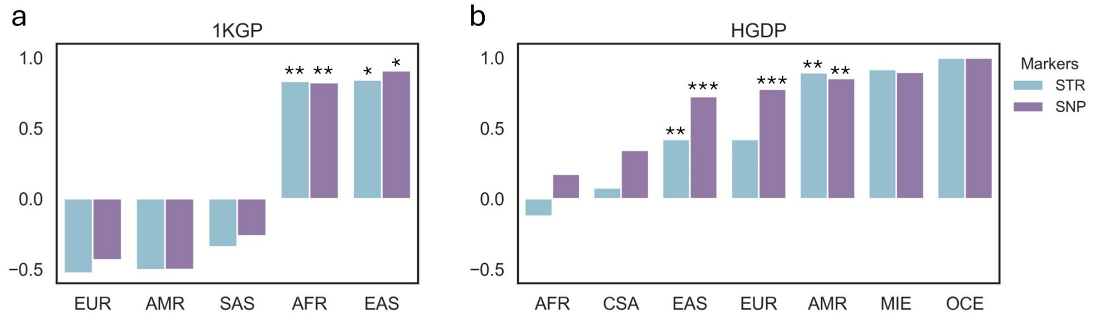


**Figure S3**: **Correlation between geographical and genetic distances across continental populations.** In each dataset, 1KGP (**a**) and HGDP (**b**), the Pearson correlation coefficients between geographical distance and genetic distance are shown for STR-based and SNP-based measures. The x-axis denotes continental populations, and the y-axis denotes the Pearson correlation coefficients. Geographical distances among regional populations were computed using the haversine formula (Earth's radius = 6,371 km). Statistical significance was assessed using a two-sided Mantel test; * $P < 0.05$, ** $P < 0.01$, *** $P < 0.001$.

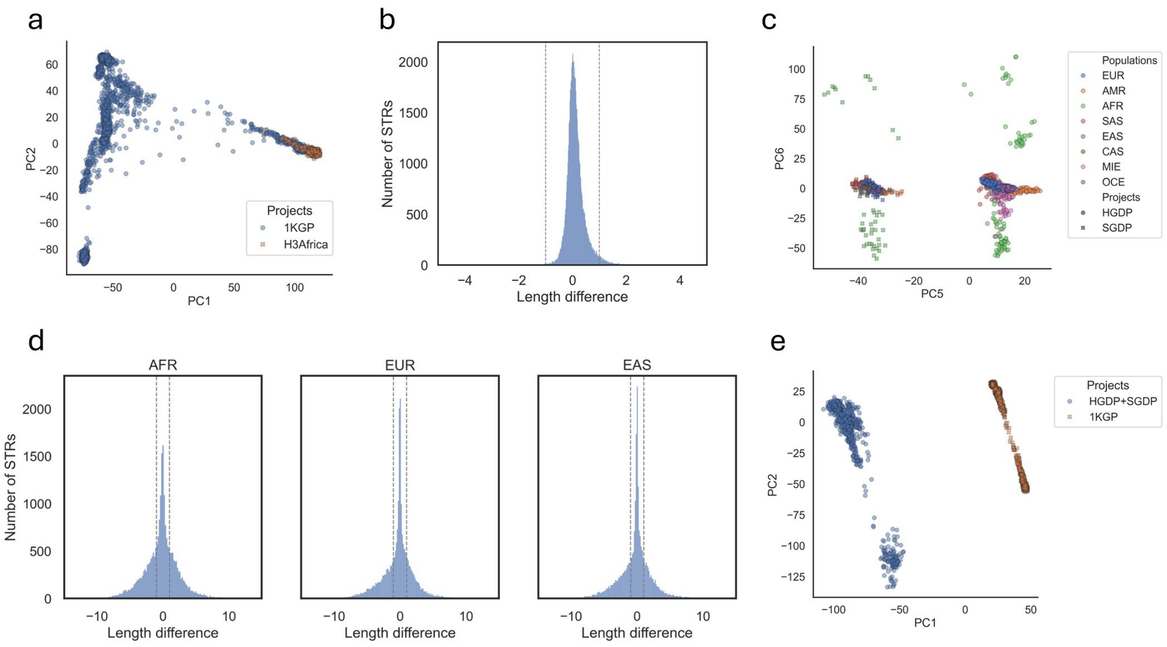


**Figure S4**: **Cross-cohort comparison of STR-based population structure.** **a** Principal component analysis (PCA) of overlapping STR loci between the 1KGP and H3Africa datasets (PC1 vs PC2) before batch effect correction. Each point represents an individual, colored and shaped by project. **a** Average STR length differences between AFR population in the 1KGP and H3Africa dataset. The x-axis denotes the average STR length differences per locus. The y-axis denotes the number of STRs. **c** PCA of overlapping STR loci between the HGDP and SGDP datasets (PC5 VS PC6). Each point represents an individual, colored by population label and shaped by project. **d** Average STR length differences between AFR, EUR, EAS populations (from left to right) from the 1KGP and HGDP+SGDP datasets. The x-axis denotes the average STR length differences per locus. The y-axis denotes the number of STRs. **e** PCA of overlapping STR loci between the 1KGP and HDGP+SGDP datasets (PC1 VS PC2). Each point represents an individual, colored and shaped by project.

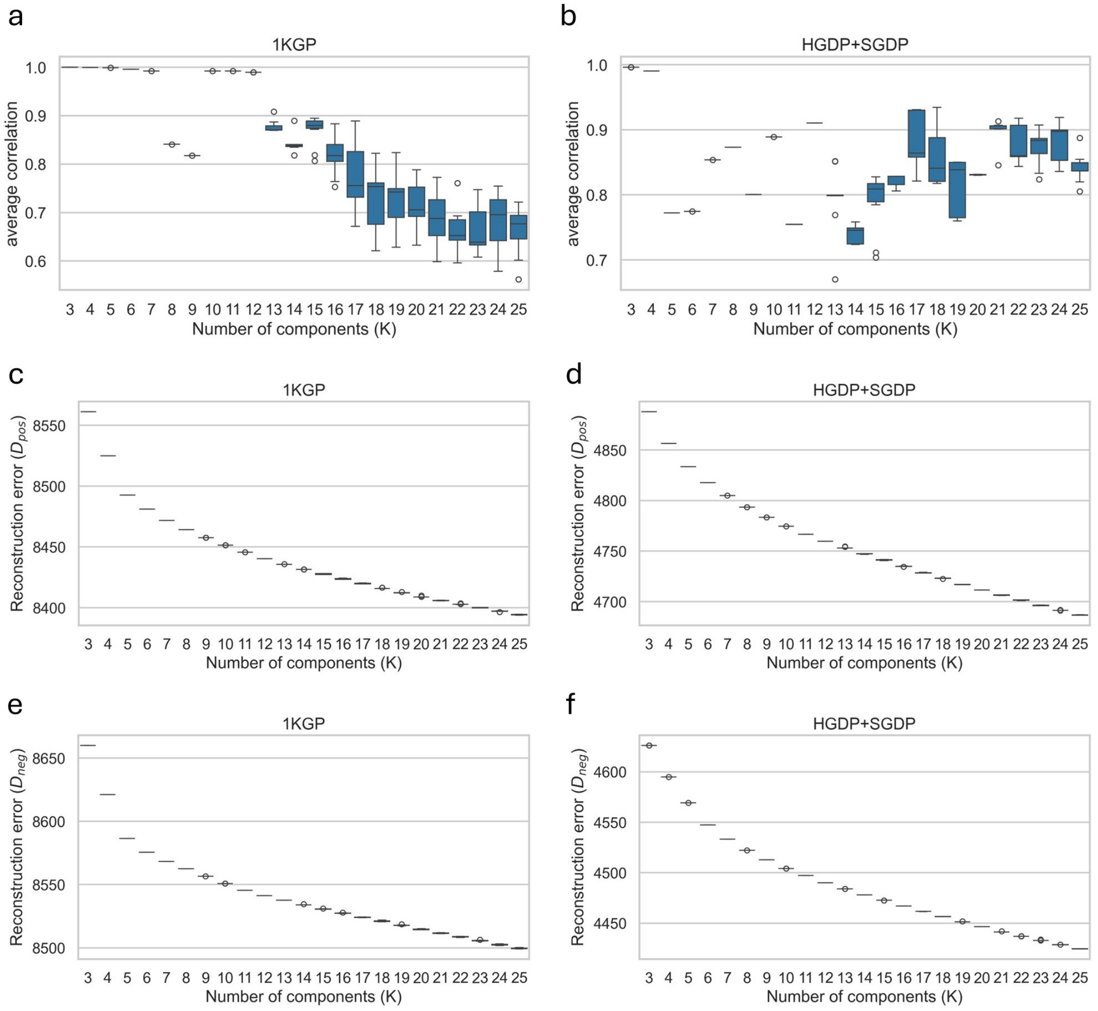


**Figure S5**: **Assessment of dNMF stability using average correlation and reconstruction error. a, b** Boxplots of average correlations across multiple runs in the 1KGP (a) and HGDP+SGDP (b) datasets. The x-axis indicates the number of components ($K$) used in the dNMF model. Each box shows the distribution of average correlation calculated from repeats runs for a given $K$. The central line indicates the median, the box boundaries represent the first and third quartiles, and whiskers extend to 1.5× the interquartile range. Outliers are shown as individual points. points. **c-f** Boxplots of reconstruction error across multiple runs for different numbers of components ($K$). Panel **c** shows results from the expansion channel in the 1KGP. Panel **e** shows results from the contraction channel in the 1KGP. **d** shows results from the expansion channel in the HGDP+SGDP. **f** shows results from the contraction channel in the HGDP+SGDP. The x-axis indicates the number of components ($K$), and the y-axis shows the reconstruction error. Each box represents the distribution of reconstruction errors across repeated runs for a given $K$.

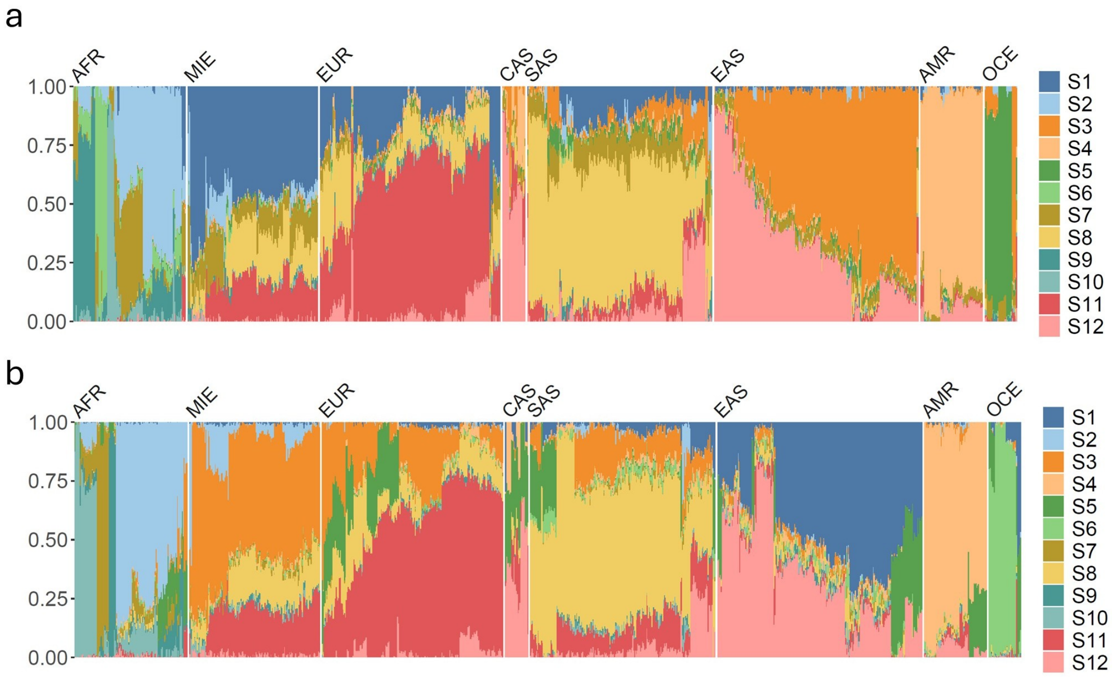


**Figure S6**: **Normalized ancestry proportion using** $K = 12$ **inferred by dNMF in the HGDP+SGDP dataset.** Panel **a** shows results from the expansion channel and panel **b** shows results from the contraction channel. Each column represents an individual, and stacked color segments indicate the proportion of each ancestry component. Individuals are ordered by hierarchical clustering. Continental population labels are shown above the plot.

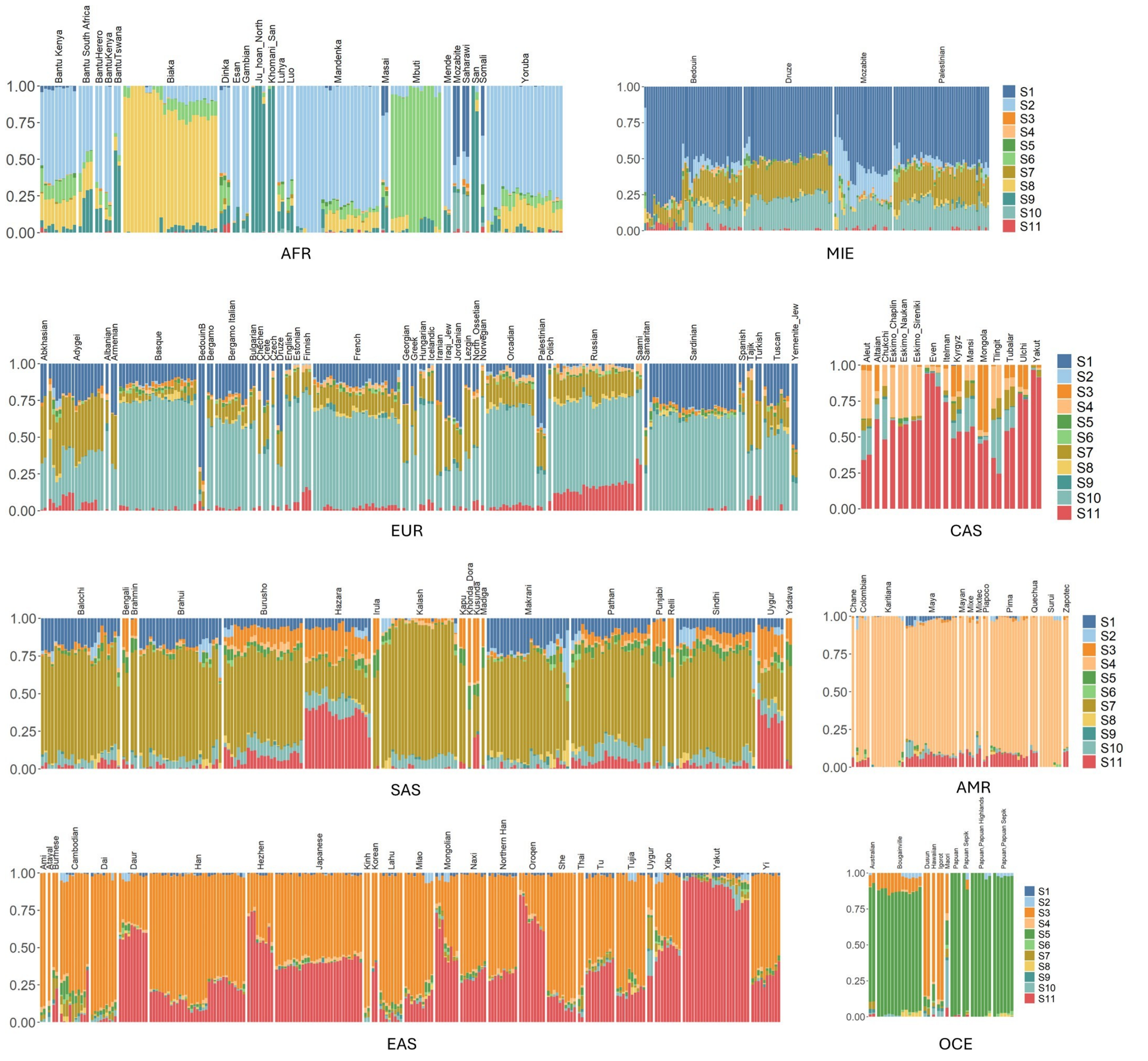


**Figure S7**: **Normalized ancestry proportion using optimal $K = 11$ inferred by dNMF in the HGDP+SGDP dataset.** Each column represents an individual, and stacked color segments indicate the proportion of each ancestry component. Regional population labels are shown above and continental population labels are shown below the plot.

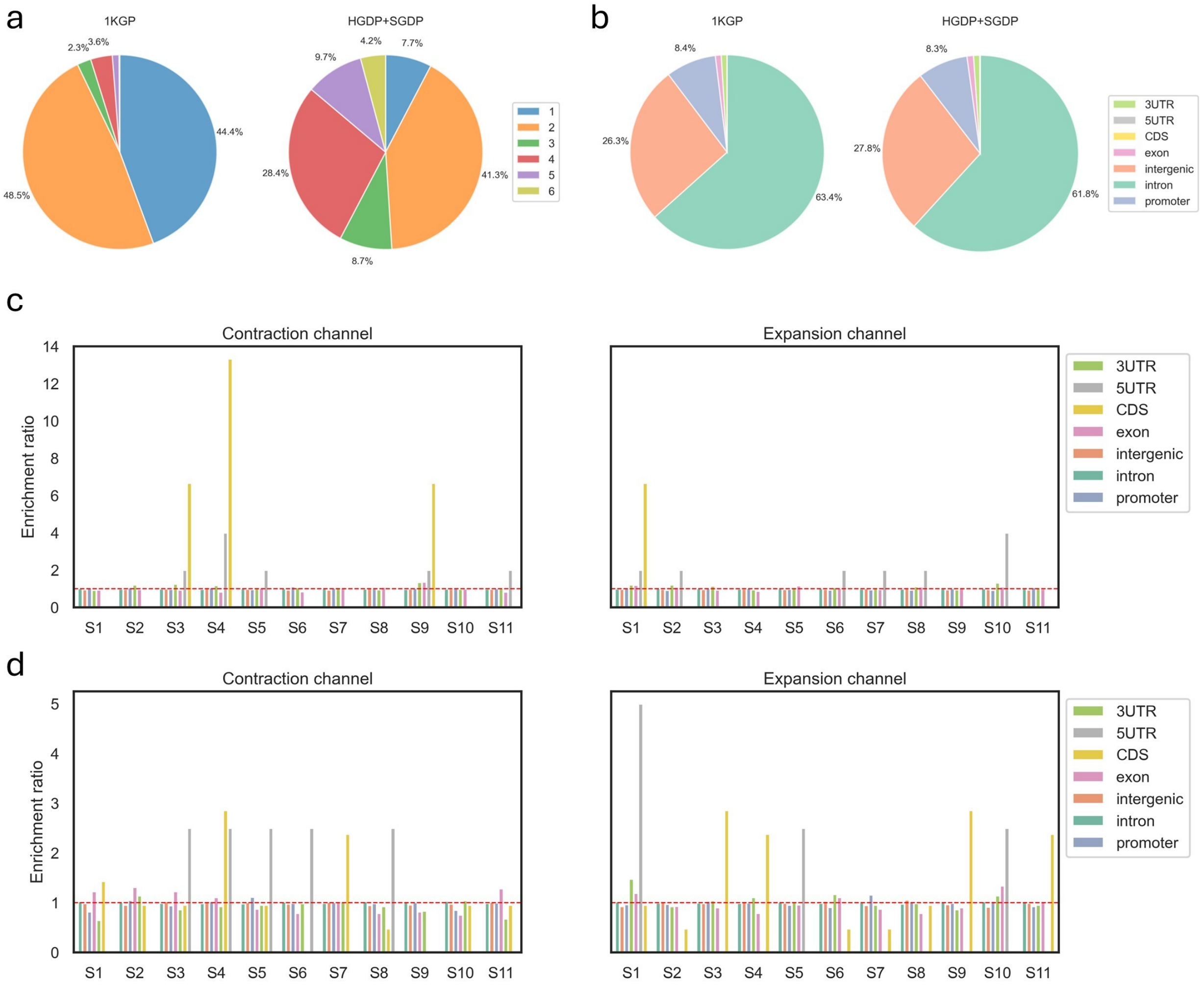


**Figure S8**: **Comparison of STR motif in the 1KGP and HGDP+SGDP dataset. a** Proportion of STR motif lengths in the input STR sets for the dNMF model from the 1KGP (left) and HGDP+SGDP (right) datasets. Each splice represents the relative fraction of STRs with a specific motif length. In the 1KGP dataset, STRs with motif lengths of 5 (1.1%) and 6 (0.1%) were omitted from the pie chart for clarity. **b** Proportion of STR located in different genomic regions in the input STR sets for the dNMF model from the 1KGP (left) and HGDP+SGDP (right) datasets. Each slice represents the relative fraction of STRs within the specified genomic category. "Exon" refers to non-coding regions of exons. **c,d** Motif genomic enrichment analysis of ancestry-informative STR signatures in the contraction (left) and expansion (right) channels for the 1KGP (c) and HGDP+SGDP (d) datasets. Enrichment ratios were computed using Fisher's exact test, with multiple testing correction performed using the Benjamini-Hochberg procedure. None of the categories were statistically significant.

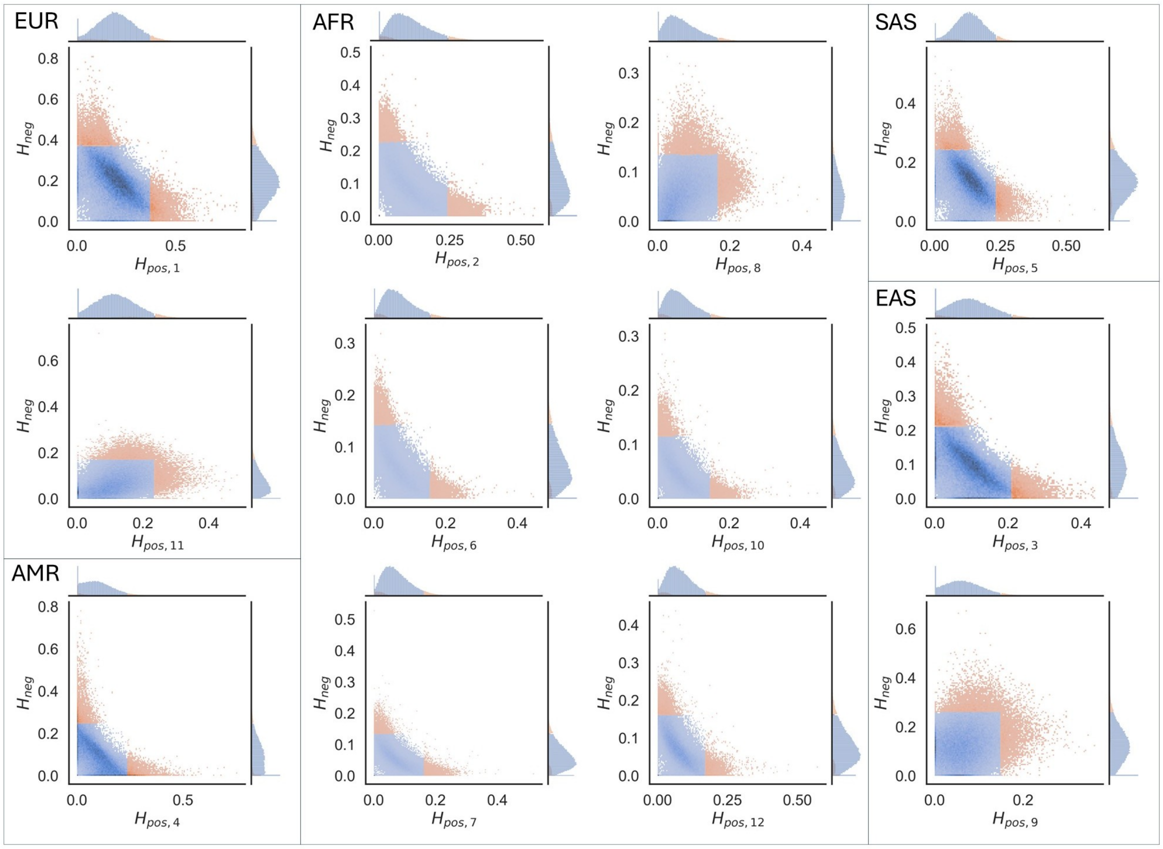


**Figure S9**: **Selection of ancestry-informative STR signatures in the 1KGP dataset.** Joint distribution of STR contribution weights between $\mathbf{H}_{pos}$ and $\mathbf{H}_{neg}$ across 12 components. In each plot, the central panel shows the two-dimensional histogram of STR contribution weights from $\mathbf{H}_{pos}$ and $\mathbf{H}_{neg}$, with marginal histograms displaying their univariate distributions. Components from $\mathbf{H}_{neg}$ are aligned to their corresponding $\mathbf{H}_{pos}$ components (see *Methods*). The top 5% of STR loci contributing most strongly to each channel are highlighted in yellow. Population labels denote the dominant population associated with each component.

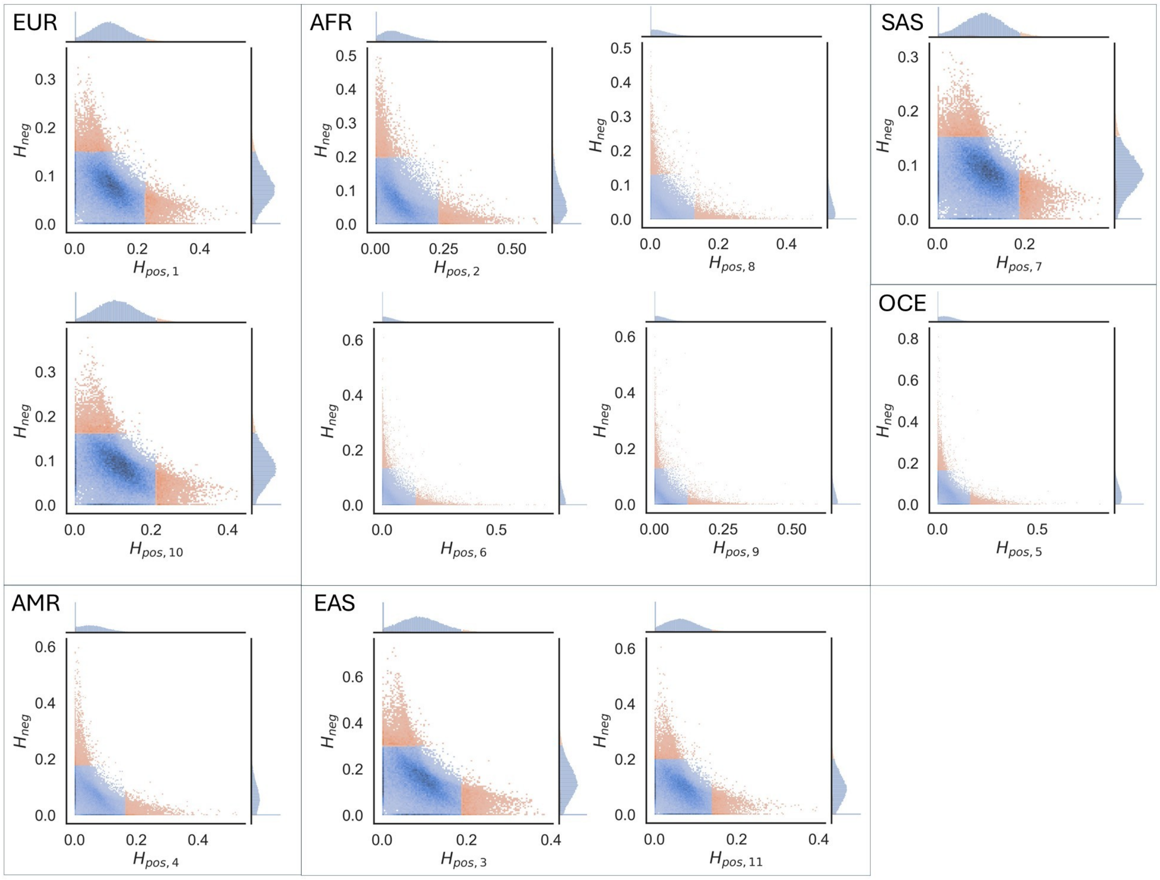


**Figure S10**: **Selection of ancestry-informative STR signatures in the HGDP+SGDP dataset.** Joint distribution of STR contribution weights between $\mathbf{H}_{pos}$ and $\mathbf{H}_{neg}$ across 11 components. In each plot, the central panel shows the two-dimensional histogram of STR contribution weights from $\mathbf{H}_{pos}$ and $\mathbf{H}_{neg}$, with marginal histograms displaying their univariate distributions. Components from $\mathbf{H}_{neg}$ are aligned to their corresponding $\mathbf{H}_{pos}$ components (see *Methods*). The top 5% of STR loci contributing most strongly to each channel are highlighted in yellow. Population labels denote the dominant population associated with each component.

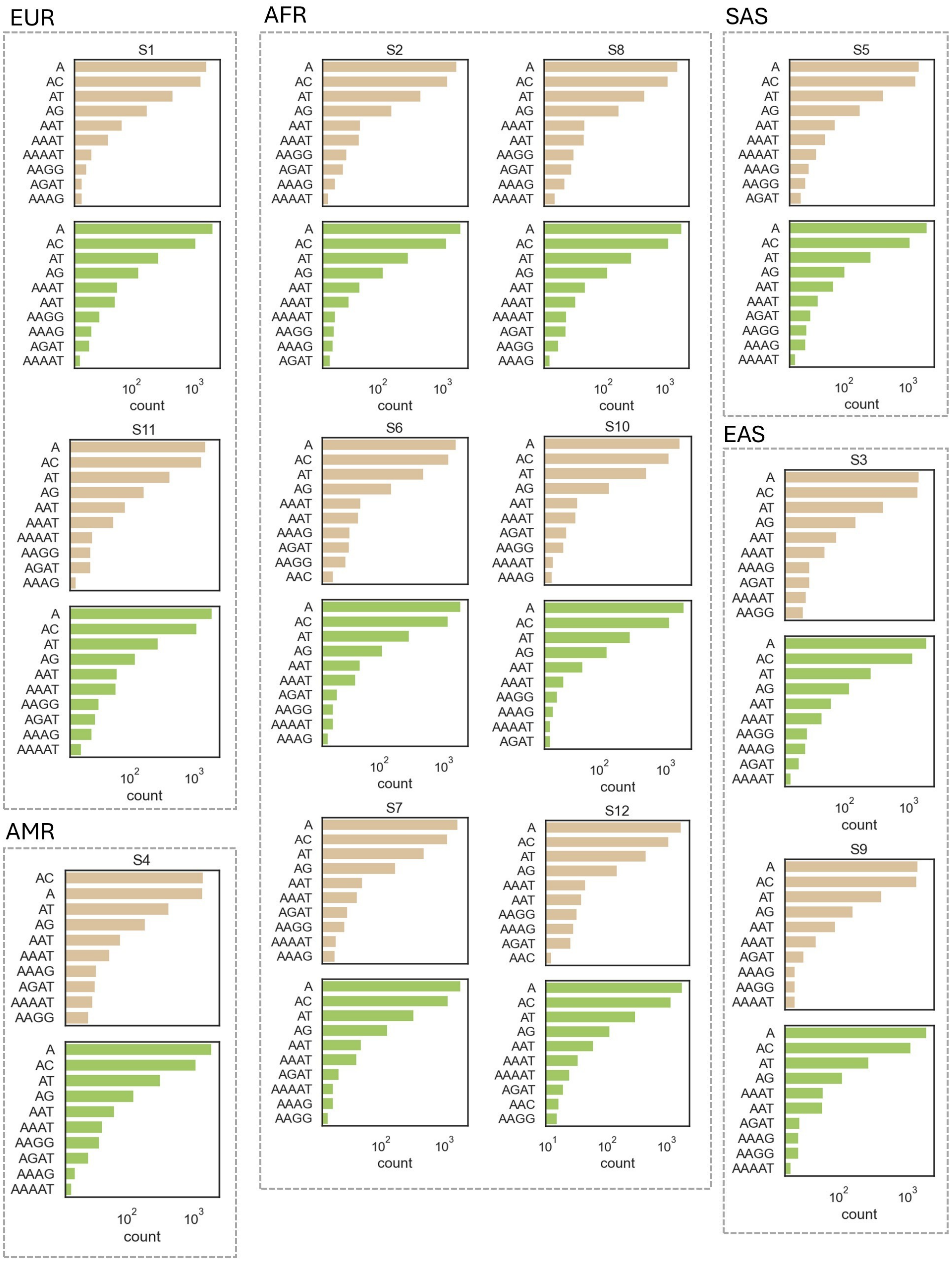


**Figure S11**: **Motif sequence composition of ancestry-informative STR signatures across components in the 1KGP dataset.** The top ten most common repeat units in each set of STR signatures are shown. STR signatures from the expansion channel are shown in yellow, and those from the contraction channel are shown in green. The x-axis indicates the number of STRs on a logarithmic scale. Population labels denote the dominant population associated with each component.

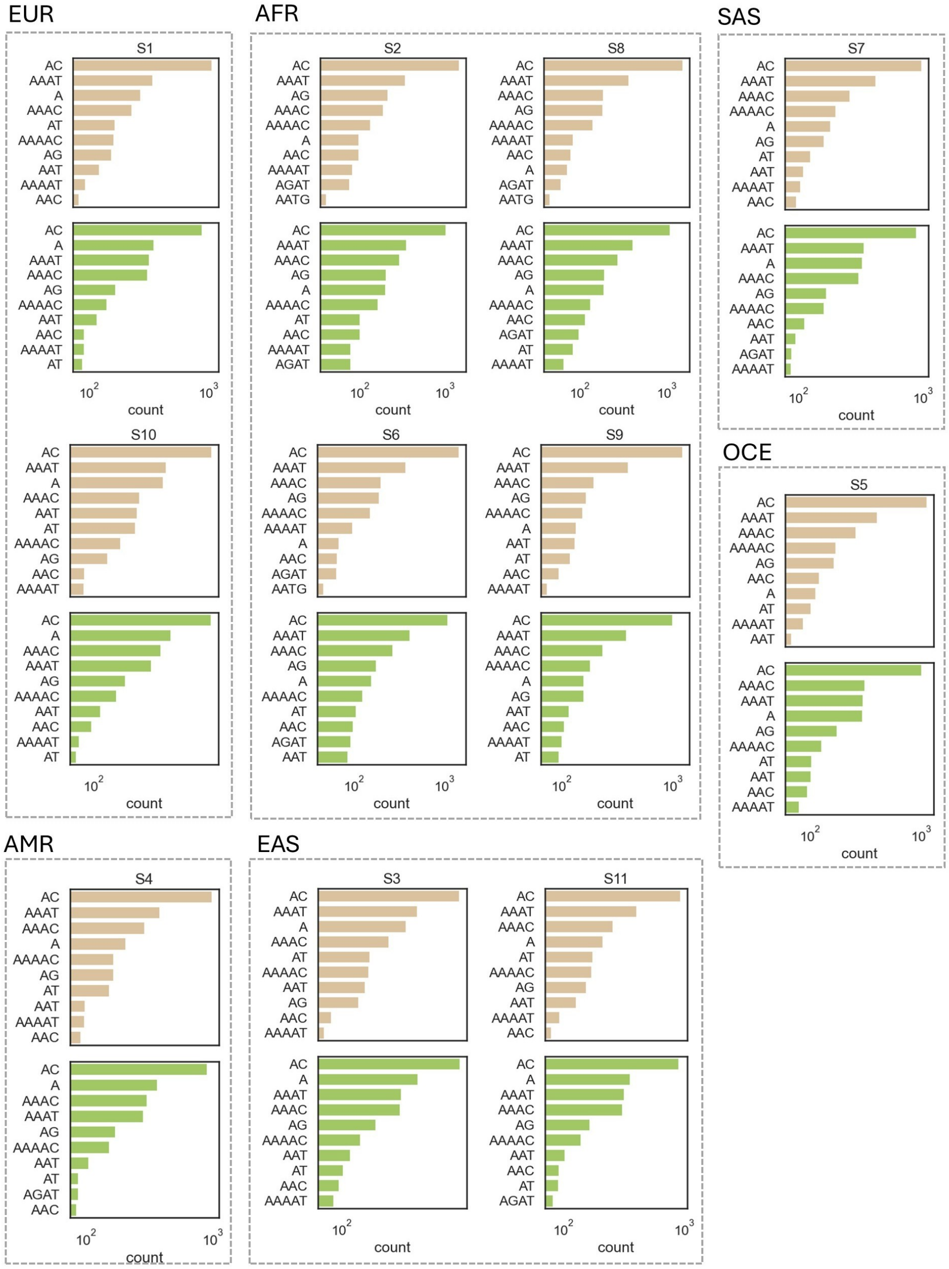


**Figure S12**: **Motif sequence composition of ancestry-informative STR signatures across components in the HGDP+SGDP dataset.** The top ten most common repeat units in each set of STR signatures are shown. STR signatures from the expansion channel are shown in yellow, and those from the contraction channel are shown in green. The x-axis indicates the number of STRs on a logarithmic scale. Population labels denote the dominant population associated with each component.

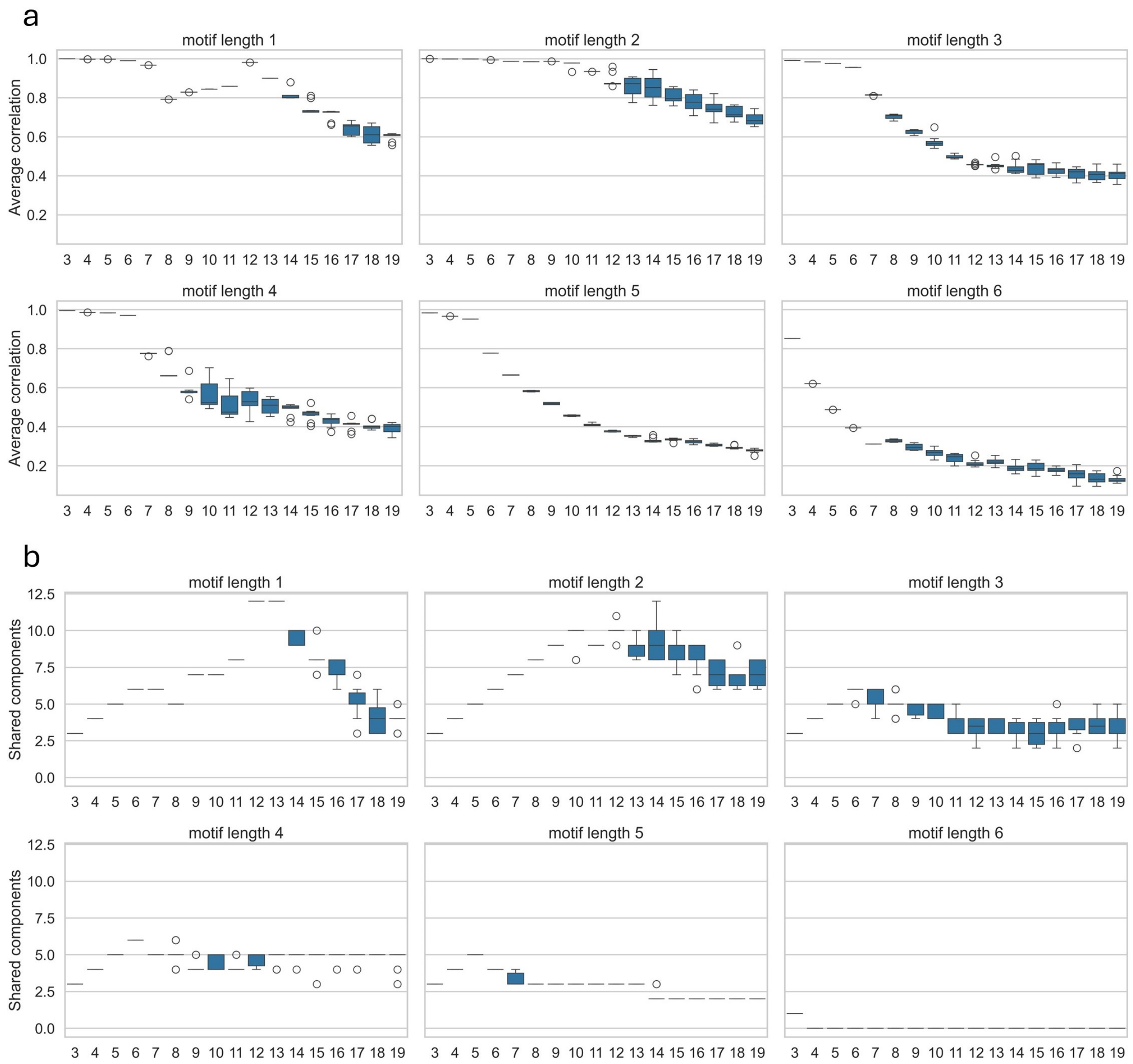


**Figure S13**: **Average correlation and shared components from motif-specific admixture inference using dNMF.** Boxplots of the average correlation (**a**) and the number of shared components (**b**) across multiple runs in the 1KGP dataset. The optimal numbers of ancestry components for motif lengths 1-5 bp are $K = 12, 10, 6, 6, 5$, respectively.
